# Nanopore-based direct RNA sequencing of the *Trypanosoma brucei* transcriptome identifies novel lncRNAs

**DOI:** 10.1101/2023.01.27.525864

**Authors:** Elisabeth Kruse, H. Ulrich Göringer

## Abstract

Trypanosomatids are single-cell eukaryotic parasites. Unlike higher eukaryotes, they control gene expression posttranscriptionally and not at the level of transcription initiation. This involves all known cellular RNA circuits, from mRNA processing to mRNA decay to translation, in addition to a large panel of RNA-interacting proteins that modulate mRNA abundance. However, other forms of gene regulation, for example, by lncRNAs, cannot be excluded. LncRNAs are poorly studied in trypanosomatids, with only a single lncRNA characterized today. Furthermore, it is not clear whether the complete inventory of trypanosomatid lncRNAs is known because of the inherent cDNA recoding and DNA amplification limitations of short-read RNA sequencing. Here we overcome these limitations by using long-read direct RNA sequencing (DRS) on nanopore arrays. We analyze the native RNA pool of the two main lifecycle stages of the African trypanosome *T. brucei* with a special emphasis on the inventory of lncRNAs. We identify 207 previously unknown lncRNAs, 109 of which are stage-specifically expressed. We also present insights into the complexity of the *T. brucei* transcriptome, including alternative transcriptional start and stop sites and potential transcript isoforms to provide a bias-free understanding of the intricate RNA landscape in *T. brucei*.

## 1. Introduction

Trypanosomatid organisms including the genera *Phytomonas, Leishmania*, and *Trypanosoma* are singlecell eukarya that parasitize plants and animals (Kaufer et al., 2020), and as such, they are of agricultural, veterinary, and medical importance. Although disease-related objectives have driven most research efforts in the past (De Rycker et al., 2023), a significant number of studies have identified a wealth of RNA-centred biochemical phenomena in the different species, including *trans-splicing*, RNA editing, ribosomal RNA fragmentation, and mitochondrial tRNA import (Clayton, 2019). This RNA-centricity is a consequence of two specific molecular characteristics of trypanosomatids. First, the organisms exhibit an unusual eukaryotic genome organization. Multiple intronless protein-coding genes are arranged in long head-to-tail tandem arrays, which are transcribed as polycistronic transcripts covering up to 100 coding sequences (CDS). Processing into monocistronic mRNAs occurs via *trans*-splicing of a capped, 39nt long spliced leader (SL)-RNA to the 5’-end and by 3’-end polyadenylation. Second, transcriptional control of gene expression is, with only a few exceptions, not existent in trypanosomatids. The organisms regulate gene expression posttranscriptionally. This implies that RNA-driven processes, such as splicing, alternative splicing, RNA turnover, and translation, must be able to sense and respond to extrinsic and intrinsic cues to adjust and modulate cellular protein levels. While a comprehensive verification of such a scenario awaits confirmation, the currently accepted view is, that changes in the transcriptome are brought about by RNA-binding proteins (RBPs) utilizing a large panel of RNA-interacting proteins (Kolev et al., 2014; Clayton, 2019). RBPs are well-documented components of the sensing and signaling pathways in higher eukaryotes and they execute their function by stabilizing or destabilizing specific mRNAs or other RNA species (reviewed in Clayton and Shapira, 2007). Trypanosomatid parasites such as the African trypanosome *Trypanosoma brucei* rely on a nearly complete inventory of canonical eukaryotic noncoding (nc)RNAs in addition to hundreds of RNA editing-specific guide (g)RNAs (Fort et al., 2022; Koslowsky et al., 2014). The panel of small ncRNAs (RNAs <200 nucleotides (nt) in length) includes the small nuclear (sn)RNA of the *trans*-spliceosome (Michaeli, 2011), the small nucleolar (sno)RNAs of the ribosomal RNA modification machinery (Martínez-Calvillo et al., 2019; Rajan et al., 2020) as well as transfer (t)RNAs and ribosomal 5S and 5.8S rRNA. Further included are cytosolic Vault (vt)RNA (Kolev et al., 2019), 7SL RNA (Michaeli et al., 1992), and telomerase RNA (Sandhu et al., 2013). Small interfering (si)RNAs are present in some but not all trypanosomatids and recent experiments identified a new class of anti-sense translational regulators, so-called TBsRNAs (Rajan et al., 2020). There is no evidence for canonical microRNAs (Clayton, 2019). Although the cellular roles for most sncRNAs are known, the situation differs drastically for long-noncoding (lnc)RNAs. LncRNAs are defined as noncoding ribonucleic acids with a nt-length >200nt. They are 5’-capped and 3’-polyadenylated and a growing number have been shown to affect transcription, DNA replication, and repair as well as cell differentiation (for a review see Statello et al., 2021). About 1500 lncRNAs are annotated in *T. brucei* (Kolev et al., 2010; Guegan et al., 2022), however, only a single lncRNA (*grumpy*) is functionally characterized. *Grumpy* acts as a precursor for the C/D box-type snoRNA *snoGrumpy*, which is involved in a vital differentiation step in the lifecycle of *T. brucei* (Guegan et al., 2022).

Although the pool of uncharacterized lncRNAs likely contains multiple regulatory lncRNAs, as well as binding targets for regulatory RBPs, it is not clear whether the annotated sequences represent the full complement of IncRNAs in *T. brucei*. This is in part due to the limited number of data but also due to the use of mainly short-read RNA sequencing (RNA-seq). Although RNA-seq has played a dominant role in transcriptome profiling in many organisms, the technique is not without limitations. Foremost, the protocol relies on reverse transcription (RT) to convert RNA into cDNA and on PCR amplification for library construction. Both enzyme treatments have been shown to introduce RT- and amplicon-specific biases (Minshal and Git, 2020), especially for sequences with low and high GC contents (Steijger et al. 2013). Other shortcomings include the misidentification of 3’-ends through internal priming events (Jan et al., 2011) and ambiguities due to RT template switching effects (Houseley and Tollervey, 2010; Mourão et al., 2019). Lastly, the requirement for fragmentation affects preferentially longer transcripts (>100nt), which makes it difficult to dissect authentic RNA processing sites from artificially introduced cleavage positions, which presents a major obstacle in quantifying and reconstructing the full sequence complexity of the RNA pool (Steijger et al., 2013).

To avoid these drawbacks, here we perform direct RNA sequencing (DRS) of the *T. brucei* transcriptome using the Oxford Nanopore Technologies (ONT) nanopore-based RNA sequencing platform (Garalde et al., 2018). ONT-DRS is a long-read sequencing technique that is capable of analyzing native RNA strands directly. The technology relies on arrays of membrane-embedded protein pores through which an electrical current is passed. Aided by a motor protein, individual RNA molecules are fed through the pores and each nucleobase can be identified by the induced shift in ion current (Garalde et al., 2018). DRS permits the analysis of full-length RNA without the need to amplify and fragment, thereby eliminating all amplification and read-length restrictions. We analyze both major lifecycle stages of the parasite (bloodstream and insect stage) and put special emphasis on cataloging all lncRNAs. We identify 207 previously unknown lncRNAs, 109 of which are expressed in a stage-specific manner. Furthermore, we provide new insights into the complexity of the *T. brucei* transcriptome including poly(A)-tail length variations, alternative splice acceptor (SAS) and polyadenylation sites (PAS), alternative transcriptional start and stop sites, and potential transcript isoforms.

## 2. Materials and Methods

### 2.1 Growth of trypanosome cells

*Trypanosoma brucei brucei* strain Lister 427 (Cross, 1975) was used for all experiments. Bloodstream-stage parasites (MITat serodeme, variant clone MITat1.2) were grown in HMI9 medium (Hirumi and Hirumi, 1989) supplemented with 10% (v/v) fetal calf serum (FCS), 0.2mM 2-mercaptoethanol and 100U/mL penicillin/streptomycin (Gibco™ Thermo Fisher Scientific). Parasites were grown at 37°C in 95% air and 5% CO_2_ at ≥95% relative humidity. Insect-stage (procyclic) parasites were propagated in SDM-79 medium (Brun and Schönenberger, 1979) in the presence of 10% (v/v) FCS at a temperature of 27°C. Parasite cell densities were determined by automated cell counting.

### 2.2. Oligodeoxynucleotide synthesis

Oligodeoxynucleotide primer molecules were synthesized by automated solid phase synthesis using 5’-Dimethoxytrityl-derivatised (β-cyanoethyl)-(N,N-diisopropyl)-phosphoramidite monomers. Full-length synthesis products were purified by reverse-phase high-performance liquid chromatography and verified by mass spectrometry. The following DNA sequences were synthesized: 18S_forward: CGGAATGGCACCACAAGAC; 18S_reverse: TGGTAAAGTTCCCCGTGTTGA; β-Tub_forward: TTCCG-CACCCTGAAACTGA; β-Tub_reverse: TGACGCCGGACACAACAG; nt_4120.1_forward: TGCAAGTGA-AGGCACTGTGT; nt_4120:1_reverse: GCAACAAGCACGGAAGATGG; nt_1759.1_forward: TTGCGTG-TGTGTTGGAATGG; nt_1759.1_reverse: ACGACTGCAAAGGAATGCCA; nt_17.3_forward: ACCCGT-AATTTGCGGACTGT; nt_17.3_reverse: TCCCTTCCGTGTATCGTTGT; nt_256.1_forward: GTGTGTAT-GGGCGTTTCGTG; nt_256.1_reverse: ACATACGCGTAGCCACACAA (18S=18S ribosomal RNA, β-Tub=β-tubulin, nt=new transcript).

### 2.3 RNA isolation

Bloodstream-stage trypanosomes were grown to 10^6^ cells/mL and insect-stage parasites to 10^7^ cells/mL. In each case, 5×10^8^ cells were pelleted at 3000xg and washed in phosphate-buffered saline (PBS: 10mM Na_2_HPO_4_, 1.8mM KH_2_PO_4_, pH7.4, 137mM NaCl, 2.7mM KCl) supplemented with 20mM glucose. Cells were shock-frozen in liquid N2, and stored at −20°C. Total RNA was isolated by guanidinium acid phenol extraction (Chomczynski and Sacchi, 1987). Polyadenylated RNA (poly(A)-RNA) was isolated from total RNA or crude cell lysates by 2 rounds of affinity chromatography using oligo d(T)_25_-derivatized magnetic beads (NEB). RNA yields were quantified by ultraviolet (UV) spectrophotometry and the integrity of the isolates was electrophoretically analyzed in 2% (w/v) agarose gels. Poly(A)-enrichment was accessed by qRT-PCR comparing the ratio of the amount of β-tubulin-specific mRNA over 18S rRNA to that of total RNA using the ΔΔCT method (Livak and Schmittgen, 2001).

### 2.4 Synthesis of cDNA and quantitative real-time (qRT)-PCR

Total RNA (200ng) or poly(A)-enriched RNA (40ng) was reverse transcribed with the help of 5U/μL Superscript IV reverse transcriptase (Thermo Scientific) in a final volume of 10μL in the presence of 2.5μM oligo dT-primer or 1μM gene-specific reverse (rev) primer. Reaction mixtures were diluted with 10-50 vol. ddH_2_O and 2μL of diluted cDNA were used in 10μL qPCR reactions containing the Luna^®^ Universal qPCR Master Mix (NEB) and 0.3μM of gene-specific forward and reverse primer. PCR reactions were carried out in a StepOnePlus real-time PCR instrument (Applied Biosystems) using a modified Fast Run protocol (5min, 95°C followed by 30 cycles of 15sec at 95°C and 30sec at 60°C). After each run, SYBR green displacement curves were generated (15sec, 95°C; 1min, 65°C followed by a 0.3°C/15sec temperature rise to 95°C) with continual fluorescence measurement at 535nm. PCR products were analyzed by electrophoresis in 10% (w/v) non-denaturing polyacrylamide gels in 90mM Tris/B(OH)3 pH8.3, 2mM EDTA buffer (TBE).

### 2.5 Preparation of DRS sequencing libraries and sequencing

Sequencing libraries were generated from 1μg poly(A)-enriched RNA following the Oxford Nanopore Technologies (ONT) SQK-RNA002 protocol. Yields were estimated fluorometrically (Qubit dsDNA-assay, ThermoFisher Scientific) and DRS sequencing was performed on a MinION device in ONT-MinION R9.4 flow cells for 20-24h. Base calling was carried out after the sequencing runs using Guppy 4.2.2 (Wick et al., 2019) as part of the ONT-MinKNOW software. Only reads with a mean *q*-score >7 were further processed.

### 2.6 Transcript mapping

A global mapping analysis of the sequenced RNAs was performed with the help of minimap2 (v2.17-r941) (Li and Durbin, 2009; Li, 2018). For this, sequencing reads were aligned to a composite reference genome consisting of the annotated genome of *T. brucei* TREU927 (v52), the mitochondrial maxicircle genome of *T. brucei* (NCBI M94286.1), and the sequence of the RNA calibration strand (RCS: yeast enolase II, YHR174W) from the Saccharomyces Genome Database (SGD). Only reads from primary alignments were further processed.

### 2.7 Identification of full-length reads

Full-length reads were identified by taking advantage of the presence of both, a 5’-spliced leader (SL)-sequence and a poly(A)-sequence at the 3’-end of every *T. brucei* mRNA. For that, sequencing reads were first mapped to the reference genome. Reads that were uniquely mapped were then appended to the 39nt SL-sequence followed by a 2nd mapping step. Reads mapping to at least 10nt of the SL-sequence were considered 5’-processed. Poly(A)-extensions were analyzed with the help of nanopolish (Loman et al., 2015; Workman et al., 2019). Only reads with both signature motifs were processed further (Supplementary Table S3).

### 2.8 Reference-free transcript identification and identification of lncRNA

Full-length reads from all libraries were combined (N=835426) and mapped to the reference genome (TriTryp v52). The alignment file was converted into a bed file using BEDTools v2.26.0 (Quinlan and Hall, 2010; Quinlan, 2014), and reads on the same strand, overlapping by at least 100nt, were clustered. Within each cluster, start and end positions were binned over a window of 100nt. Within each bin, the most frequent positions were considered transcript boundaries. Exons supported by at least 3 reads (N=10652 out of a total of 18887 exons/transcript variants) were considered for the subsequent analysis. lncRNAs were identified from the pool of ≥200nt transcripts (supported by minimally 3 reads) using CPC2 (Kang et al., 2017) and Lncfinder (Han et al., 2019).

### 2.9 Estimation of transcript abundance, DGE analysis, and statistics

Expression profiles and the analysis of differential gene expression (DGE) were performed using the Bioconductor packages RsubRead (Liao et al., 2019) and edgeR (Robinson et al., 2010). *p*-values were adjusted according to Benjamin and Hochberg, and statistical calculations were performed using R (version 4.0.3) and Rstudio (version 1.1.453).

## 3. Results and Discussion

### 3.1 RNA sequencing libraries - quality assessment

We prepared three libraries for each, the bloodstream and insect life cycle stage of the parasite. Per library, we obtained between 2.8×10^5^ and 6.6 x10^5^ reads (Supplementary Table S1), and approximately 90% of all reads were base-called with a median Phred quality (*q*)-score ≥7. Both the read quality and the read lengths are comparable for the bloodstream and insect stage libraries with a median length of 770±57nt, a median *q*-score of 10.2±0.3, and *p*-values of 0.8 and 0.43. Between 92-98% of the reads were mapped to the reference genome of *T. brucei* TREU927 and the mitochondrial (maxicircle) genome of the parasite. The mapping frequencies (Supplementary Table S2) are lower for libraries from bloodstream-stage parasites (*p*=1.5×10^-4^), which is due to the high level of variant surface glycoprotein (VSG) expression. Three to four percent of all reads map to this transcript. Since the specific VSG sequence is lacking in the annotated TREU927 genome, a higher number of unmapped reads is obtained. Contaminations with ribosomal RNAs are <1.4% and the genome coverage varies between 64% and 73%. 79-84% of the annotated exons (N=10728) are detected, with significantly more exons covered in the bloodstream-stage libraries (*p*=0.0014). As mentioned above, this is due to the presence of reads for VSG and expression site-associated gene (ESAG) transcripts. The overall error frequency for all libraries enumerates to about 8%. Gene expression profiles within libraries from either developmental stage correlate with a Spearman rank correlation coefficient (*ρ*) of 0.9. Procyclic and bloodstream-stage libraries correlate with *ρ*=0.75.

### 3.2 Identification of full-length transcripts

In contrast to short-read RNA-seq, which requires the assembly of overlapping reads, nanopore-based direct RNA sequencing (DRS) monitors the transcript inventory of cells “directly”. This enables a straightforward mapping of gene boundaries and transcript termini, which are characterized by sharp changes in the genome coverage profiles. As a consequence, DRS allows the identification and perhaps the correction of transcript termini, and it enables the detection of transcript isoforms of novel, nonannotated transcripts, and RNA maturation intermediates. Examples for each case are shown in Figure 1. Intermediates resulting from endonucleolytic cleavage of precursor RNAs such as snoRNA or rRNA biogenesis products also produce discrete steps in the coverage plots but lack spliced leader (SL) and poly(A)-tail signatures (Figure 1c,d).

**Figure 1.**
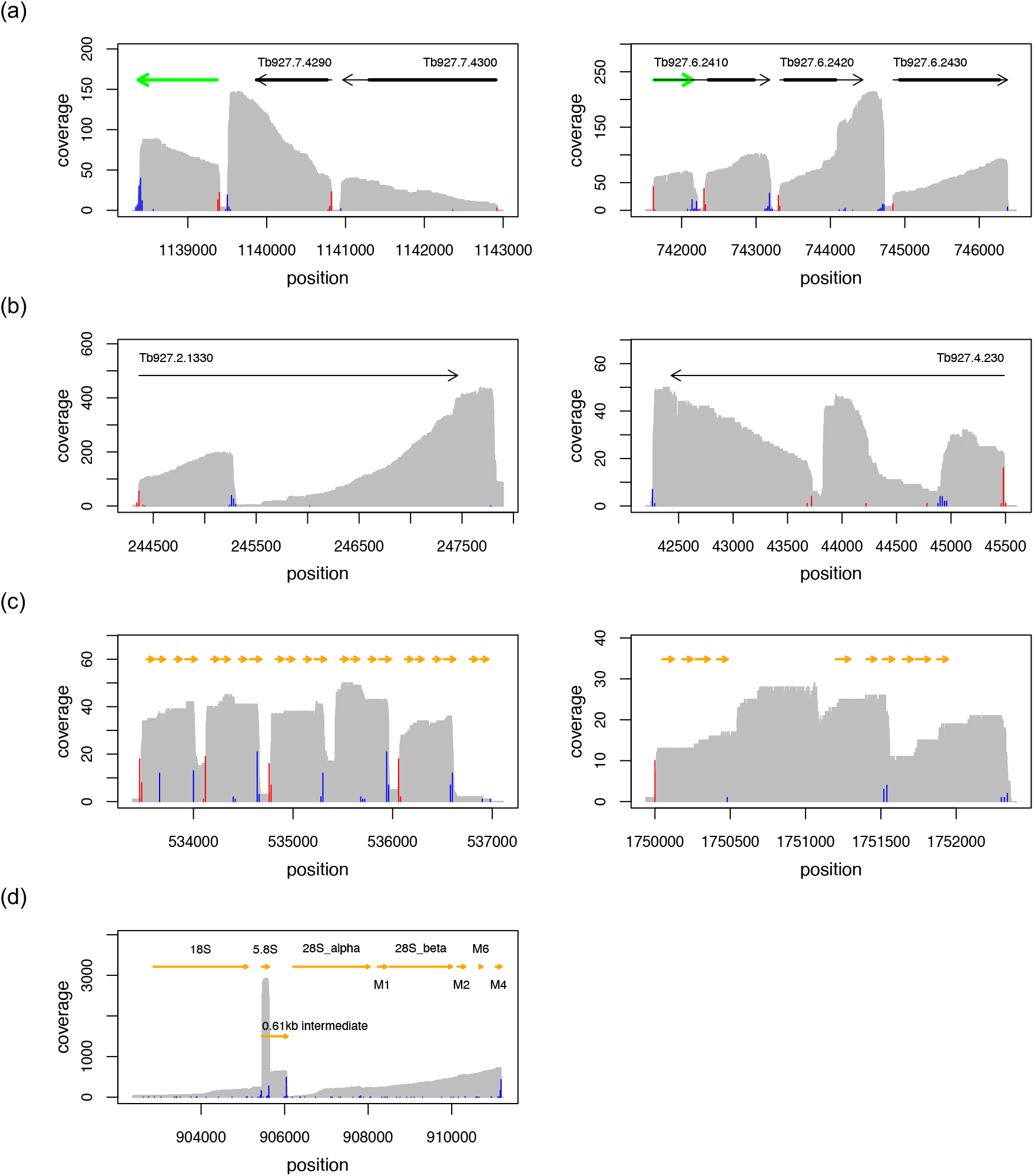
Coverage plots (grey) of representative nanopore-based direct RNA sequencing results. (a) Precise mapping of annotated transcript boundaries, correction of annotated UTRs, and detection of novel intergenic (left) or UTR-based transcripts (right). Grey arrows are annotated genes and thick, black lines denote coding sequences. Green arrows are novel transcripts. Splice acceptor sites are in red, and polyadenylation sites are in blue. (b) Mapping of pseudogene-derived transcripts. (c) Mapping of snoRNA precursors (arrows in yellow). (d) Detection of rRNA maturation intermediates (arrows in orange). The data in (a) and (b) are derived from bloodstream-stage trypanosomes. The data in (c) and (d) are from procyclic-stage parasites. The following genomic regions are presented: (a) Tb927_07_v5.1:1138300-1143000 and Tb927_06_v5.1:741520-746500. (b) Tb927_02_v5.1: 244300-247900 and Tb927_04_v5.1:42200-45600. (c) Tb927_08_v5.1:533390-537110 and Tb927_10_v5.1: 17499 40-1752400. (d) Tb927_03_v5.1:902360-911230.

In living cells, full-length transcripts coexist with RNA degradation products, which result from cotranslational mRNA decay, nonsense-mediated mRNA decay, or nucleolytic RNA maturation processes. Since DRS starts from the 3’-end of poly(A)-tailed RNAs, this results in a 3’-end bias. Furthermore, DRS also suffers from a 5’-end ambiguity because the sequenced RNA strands are released from the nanopore about 10–12nt before they reach the 5’-terminus (Jain et al., 2022). Artificial 5’- and 3’-termini are randomly distributed and generally represented by only single reads. By contrast, *true* transcript boundaries show higher read counts and are clustered toward the start and end of a transcript (Supplementary Figure S2).

As stated above, translatable mRNAs and lncRNAs in trypanosomatids are generated by *trans*-splicing and polyadenylation from polycistronic primary transcripts. Thus, mature transcripts are represented as polyadenylated reads with a spliced leader (SL)-sequence at the 5’-end. This unique feature was used to unambiguously identify full-length transcripts. Reads carrying the SL sequence were selected as detailed in the Materials and Methods section. The error rate of this approach was estimated from the number of putative SL-containing reads for the spike-in control transcript, which is <1%. Due to the 3’-bias, less than 60% of all reads exhibit the 5’-SL-sequence, with significantly fewer 5’-full-length reads (27-37%, *p*=0.005) in the libraries derived from bloodstream-stage parasites (Supplementary Table S3). *In toto*, we obtained 9.7 x10^5^ reads containing parts of the SL-sequence representing 86686 putative splice acceptor sites. 80% of the reads that mapped to annotated exons are localized in the 5’-UTRs, but about 10% map within coding sequences. These may represent transcript isoforms that encode truncated proteins, novel intragenic lncRNAs, or perhaps reflect misannotations.

Homopolymers in the sequence are characterized as stretches of low variance in the raw DRS current signal. Homopolymer signals immediately following the signal of the adapter sequence are indicative of a poly(A)-tail, and the extent of this signal can be used to calculate the tail length (Krause et al., 2019, Workman, 2019). We used the nanopolish poly(A)-module (Loman et al., 2015; Workman et al., 2019) to identify reads containing poly(A)-tails and to access their length. Altogether, 84% of all reads (from all libraries) were identified as polyadenylated (Supplementary Table S3). 83% of the reads for the spike-in control and 60% of the reads mapping to the mitochondrial genome were polyadenylated. Also, 83% of the reads mapping to rRNA genes were polyadenylated. The polyadenylated reads could be assigned to 323153 unique positions in the *T. brucei* genome. About 40% of the polyadenylation sites were assigned to annotated exons, the majority mapping to 3’ UTRs. The median poly(A)-tail length is 96nt for annotated mRNAs. It is moderately higher for pseudogenic transcripts and lncRNAs (117nt), but considerably shorter for rRNAs (26nt) and mitochondrial transcripts (29nt) (Figure 2). For the spike-in control, the median tail length matches the reported value of 35nt (Maier et al., 2020). Note that the number of putative polyadenylation or splice acceptor sites is likely overestimated. The inherent error rate of nanopore sequencing skews the identification of the exact position of polyadenylation sites. Indels immediately upstream of the poly(A)-tails or downstream of putative SAS sites will cause a deviation from the true sites and also lead to a lower read count at the corresponding positions. However, globally the DRS data agree well with the results obtained from short-read RNA-seq (for a comparison see Supplementary Figure S4).

**Figure 2.**
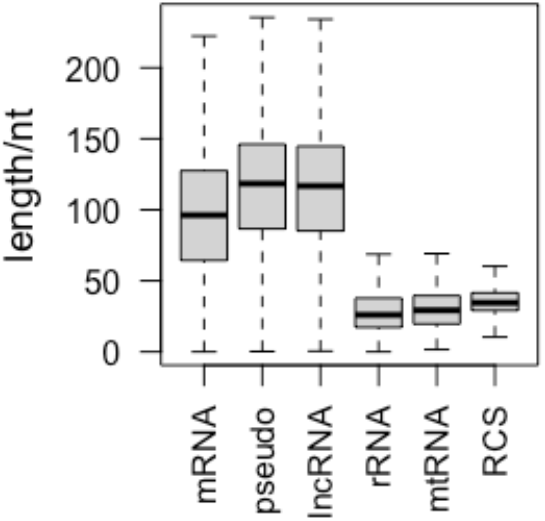
Poly(A)-tail length distribution for different transcript classes. Transcripts are grouped based on the annotated features in TriTrypDB-52_TbruceiTREU927.gff and the published lncRNAs in Guegan et al., 2022. RCS: RNA-Calibrant-Strand used in the library preparation (yeast ENO2). Bold horizontal lines represent the median.

### 3.3 Reference-free transcript identification

The selection of full-length reads with both, a 5’-SL and a 3’-poly(A)-sequence on the same transcript, enabled us to precisely map all splice variants or transcript isoforms thereby improving the current transcriptome annotation. For that, we used all full-length reads from all libraries (N=832091) and identified 18406 exons and/or transcript isoforms, many of which are partially overlapping. 30% of the transcripts/isoforms consist of only 1 read. The maximum number of reads for a single transcript is 6076, which encodes the ribosomal protein L11 (Tb927.9.7620). The transcript length distribution varies from 113nt to 14636nt with a median of 1501nt. The longest transcript (14636nt) covers the complete coding sequence of the trypanosome axoneme component hydin (Dawe et al., 2007) (Tb927.6.3150). Four other transcripts >10000nt contain the complete coding sequences of the paraflagellar rod component 2, the ankyrin-repeat protein, the flagellum attachment zone protein, and the neurobeachin/beige protein (Tb927.6.3670, Tb927.9.15400, Tb927.9.2075, Tb927.10.6150) and comprise 3’- and/or 5’-extensions indicative of incompletely annotated UTRs. All transcripts >200nt with more than two reads (N=10840) were considered for subsequent analysis. These transcripts/isoforms were assigned to 7393 loci or clusters containing up to 17 different exons or isoforms per transcript. About 50% (N=5637) of these clusters represent a single exon, while 1756 loci encode 5203 different transcripts/isoforms in total. They either represent splice variants of the same transcription unit or result from alternative or aberrant splicing events thereby giving rise to split or truncated exons or to di- or poly-cistronic transcripts.

### 3.4 Comparison to annotated exons

Of all transcripts with a nucleotide length >200nt and supported by >2 reads, 829 do not overlap with any annotated exon. 72 overlap by <10% of their nucleotide length. Filtering out exons overlapping with the lncRNAs reported by Guegan et al., 2022 resulted in 538 novel transcripts. However, this approach underestimates the true number since splice-variants leading to split or truncated mRNAs in addition to pseudogene-derived transcripts are not considered. The annotated *T. brucei* chromosomes contain 9687 cistrons that are expected to be *trans*-spliced and polyadenylated. Of these, 7358 are at least partially covered in our DRS data. Taking into account an uncertainty of ±20nt in the mapping of transcript boundaries, 2319 observed transcripts match precisely their annotated counterpart. About 10% of the sequenced transcripts represent partial sequences of annotated exons. More than 25% of the annotated exons are completely covered and include the 5’- and 3’-extensions (median length=704nt), which is indicative of incompletely annotated UTRs. Large discrepancies between the annotated and observed length of transcripts can also be attributed to splicing events, which result in truncated RNAs, or to aberrant *trans*-splicing and polyadenylation events that generate long di- or poly-cistronic transcripts. Based on the current annotation, 400 observed exons completely cover two annotated exons, and thus, are di-cistronic. Sixteen transcripts cover three annotated exons. Small ncRNAs were omitted from the analysis since many of them share a common precursor and thus, *per se*, will cover multiple annotated exons. Transcripts covering >1 exon may be the result of aberrant *trans*-splicing or polyadenylation or simply the result of misannotation. However, in some cases, di-cistronic transcripts account for >20% of all sequencing reads, which argues against a *trans*-splicing or polyadenylation error.

### 3.5 Long noncoding RNAs

To revise and potentially expand the inventory of lncRNA in *T. brucei* we analyzed all identified full-length transcripts for their coding potential. Two prediction tools that depend on different search criteria were used: CPC2 makes use of the Ficket score, the integrity of the open reading frames, and includes physicochemical properties of sequence-derived peptides (Fickett and Tung, 1992). The algorithm is claimed to be species-neutral (Kong et al., 2007, Kang et al., 2017). Lnc_finder from the R package LncFinder (Han et al., 2019) relies on intrinsic features of nucleic acid sequences and can be “trained” for any organism of interest. Of 10652 sequences, 2445 were predicted to be noncoding by CPC2. LncFinder, trained on the TriTrypDBv52 *T. brucei* sequences, identified 1806 putative lncRNAs, and the sequences predicted by both methods (N=1801) were used for subsequent analysis. Interestingly, both search algorithms also predicted annotated protein-coding sequences as noncoding (17% for CPC2, and 5% for LncFinder, Supplementary Table S4).

#### 3.5.1 Characterisation of long noncoding RNAs

The predicted lncRNA genes cluster at 1382 genomic loci covering all 11 Mbp-size *T. brucei* chromosomes (Figure 3a,b and Supplementary Figure S5). The 4 annotated small chromosomes of *T. brucei* code for an additional 4 lnc-transcripts. 1139 loci are represented by a single transcript. Of the remaining 243 clusters, 60% contain 2 and 40% up to 17 transcripts, which represent splice variants of the same gene and/or overlapping transcripts coded by adjacent loci (Figure 3c,d). 62 sequences cover annotated snoRNAs. These transcripts are located at 27 loci with 12 loci coding for overlapping transcripts (Supplementary Figure S6). Since snoRNA precursors do not represent functional lncRNAs, these sequences were omitted from the analysis, resulting in 1739 transcripts in 1355 clusters. Based on the current annotation of the *T. brucei* transcriptome (TriTrypDBv52), 748 putative lncRNAs (43%), clustering at 585 loci, do not overlap with any annotated features, and thus formally represent intergenic sequences. However, since the UTR annotation in the reference genome is incomplete, the true number of intergenic lncRNAs is likely lower. Using the information for transcripts identified from full-length reads, only 599 transcripts (at 452 loci) are identified as intergenic. 153 transcripts in 133 clusters partially overlap with coding sequences of annotated mRNAs. 80 transcripts in 54 clusters map to annotated pseudogenic transcripts, and thus might represent pseudogene-derived lncRNAs. Of the 1491 annotated lncRNAs in the reference genome, 50% are confirmed by our DRS data. 27% overlap by at least 10% of their nt-length and about 20% of the annotated lncRNAs are not detected as independent transcripts but as part of UTRs.

**Figure 3.**
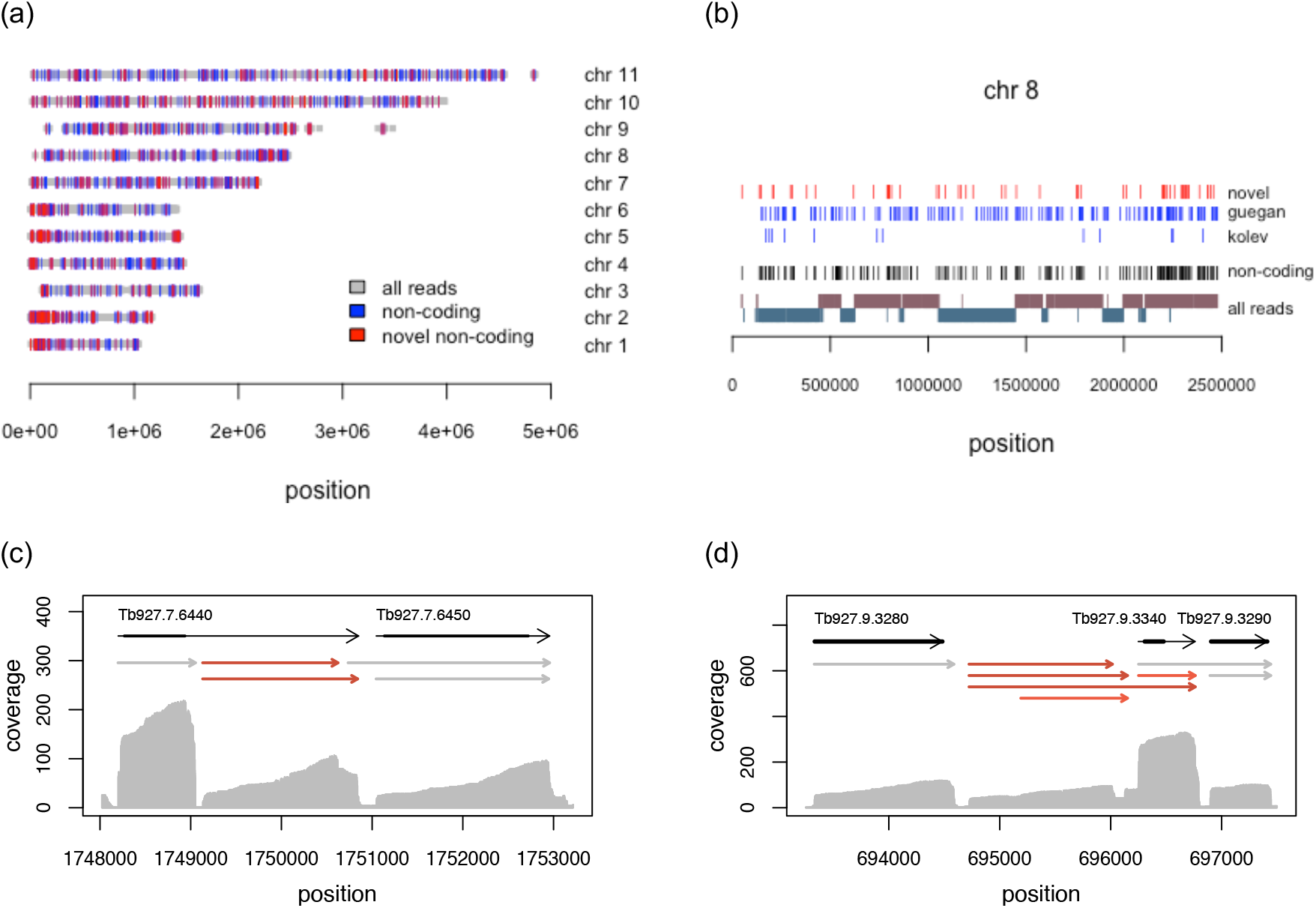
Genomic localization of lncRNA genes. (a) Localization of lncRNAs genes on the 11 large (Mbp) chromosomes of *T. brucei*. Regions covered by reads, based on all bloodstream- and procyclic-stage libraries are in grey. Known lncRNA genes (Kolev et al., 2010; Guegan et al., 2022) are in blue. Novel lncRNA genes (this study) are in red. (b) Detailed view of chromosome 8. The localization of all novel lncRNA genes (red) is separated from the published sequences of Kolev et al., 2010, and Guegan et al., 2022 (blue). For comparison, “all reads” are shown in a strand-specific manner. The top line represents the plus strand, and the bottom line the minus strand. (c) Clustering of new putative lncRNAs (red) as “minor” splice acceptor or polyadenylation site variants. (d) Complex clustering of novel lncRNAs. Coverage plots (grey) from the combined reads of all bloodstream-stage libraries are shown. Grey arrows represent annotated transcripts with coding regions as thick black lines. Grey arrows are predicted coding transcripts and red arrows represent new lncRNAs.

A general comparison of the coding and noncoding transcripts of *T. brucei* for a set of sequence and structure-specific parameters is provided in Supplementary Figure S7. This includes the nt-length distribution, the nucleotide and dinucleotide content of the RNAs, as well as the thermodynamic stability (ΔG) of their 2D minimum free energy fold (MFE). With a median of 750nt, the pool of noncoding RNAs is shorter than that of coding RNAs (median=1600nt). The molecules have a lower GC content (0.45 *versus* 0.49) and fold into less stable 2D structures (ΔG=-0.28kcal/mol/nt *versus* −0.32kcal/mol/nt) (Supplementary Figure S7).

#### 3.5.2 Novel lncRNAs

Removing from the pool of intergenic noncoding transcripts all annotated lncRNAs resulted in 207 novel intergenic loci: 251 transcripts with up to 6 transcripts per cluster. Examples are shown in Supplementary Figure S8. Some co-localize with transcription start sites and/or loci coding for variant surface glycoproteins (VSG) or expression site-associated genes (ESAG) (Figure 4). The sequences from 41 loci are not unique with up to 4 copies and some have additional counterparts in the genome, which are either not expressed or not detected. These redundant sequences may be arranged as repeats of alternating lncRNA and genes that encode the same protein. Examples are shown in Figure 5. A list of all newly identified lncRNA is provided in Supplementary Table S5.

**Figure 4.**
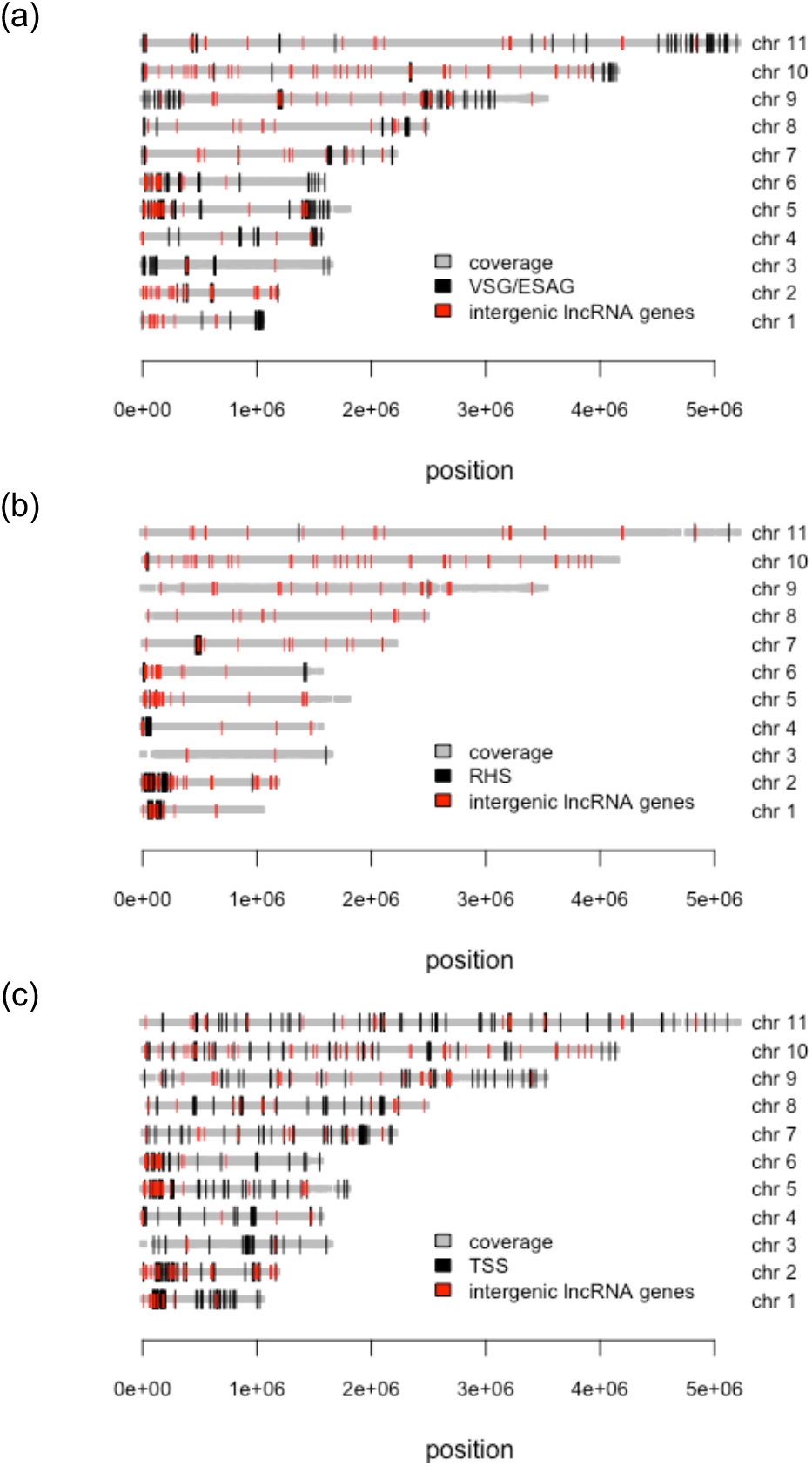
Localization of novel intergenic lncRNA genes (red) on the 11 Mbp chromosomes (chr) of *T. brucei*. (a) Comparison to the positions of variant surface glycoprotein (VSG) genes and expression site-associated genes (ESAG). (b) Comparison to the locations of genes of the retrotransposon hot spot gene family (RHS). (c) Comparison to transcription start sites (TSS). Bold grey lines: chromosomal regions covered by sequencing reads.

**Figure 5.**
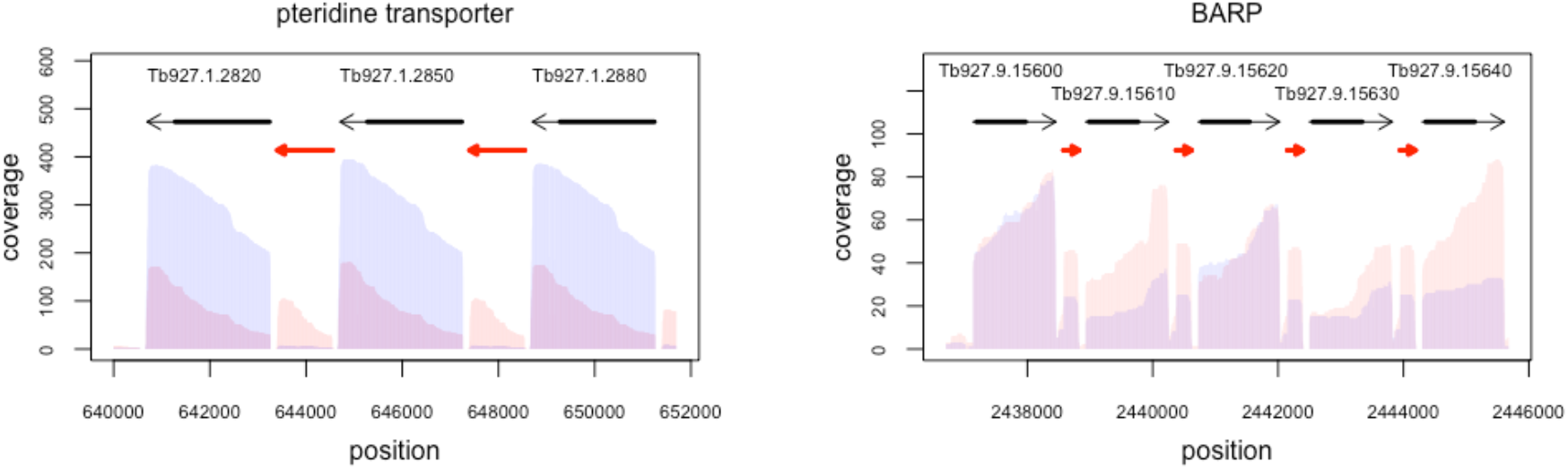
Repetitive lncRNAs. Two examples of genomic regions consisting of repeats of protein-coding genes for identical gene products interrupted by identical lncRNAs. Left: Tandem repeat of pteridine transporter genes on chromosome 1. Right: 3’-domain of the tandem gene array of the bloodstream-alanine-rich-protein (BARP) on chromosome 9. Coverage profiles from bloodstream-stage *T. brucei* are in red. Procyclic-stage profiles are in blue. Grey arrows are annotated exons with coding sequences as thick black lines. Red arrows are lncRNAs.

### 3.6 Differential gene expression (DGE) between insect- and bloodstream-stage trypanosomes

Short-read RNA-seq has been widely used to analyze changes in gene expression during the *T. brucei* life cycle (Veitch et al., 2010, Siegel et al., 2010, Christiano et al, 2017). However, the requirement for reverse transcription and PCR during library preparation has been shown to skew the quantification of transcript levels (Aird et al., 2011, Parekh et al., 2016). By contrast, DRS has been shown to accurately enumerate transcript amounts (Byrne et al., 2017, Jenjaroenpun et al., 2018, Gleeson, 2022), which tempted us to compare the DRS-derived transcript levels between bloodstream- and insect-stage trypanosomes. Genes showing at least a 4-fold change in transcript level with adjusted *p*-values <0.05 were considered differentially expressed. Of 11769 genes, 8355 were included in the analysis based on the criterion of ≥15 reads/gene in all libraries and ≥5 reads in either the procyclic- or bloodstream-stage libraries. 417 genes are up-regulated in bloodstream-stage parasites compared to procyclic trypanosomes, and 155 genes are down-regulated. As expected and shown in Figure 6, the transcripts for stage-specific surface glycoproteins VSG, ISG, and procyclin, as well as expression site-associated genes (ESAG, PAG) are over-represented in the panel of differentially expressed genes (*p*=4.5×10^-83^ Fisher’s exact test) and the same holds for members of the retrotransposon hotspot protein multigene family (RHS). This is consistent with the observation that many RHS genes or RHS pseudogenes are associated with VSG and ESAG loci in sub-telomeric regions (Bringaud et al., 2002, Bringaud et al., 2004). A list of all differentially expressed genes is provided in Supplementary Tables S6 and S7.

**Figure 6.**
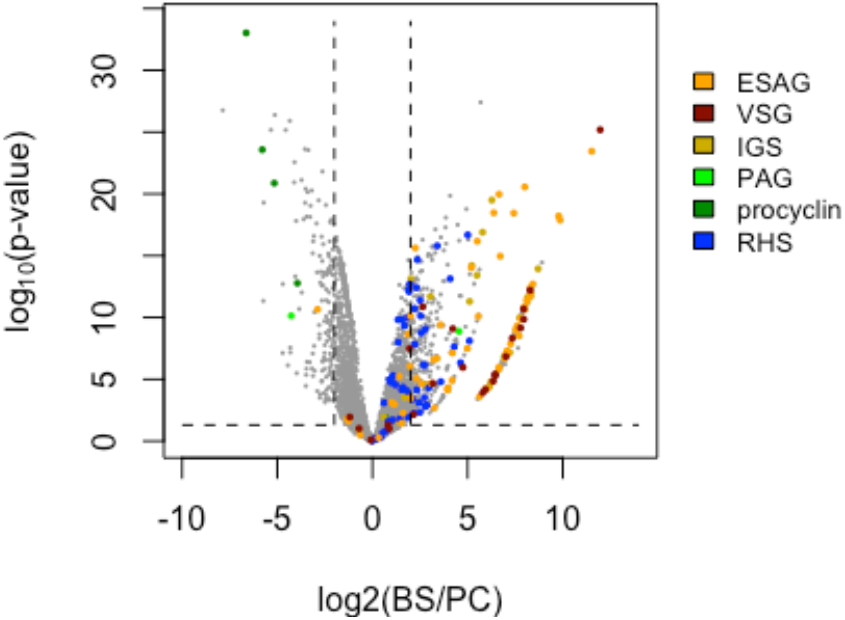
Volcano plot representation of differential gene expression between bloodstream-stage (BS) and procyclic-stage (PC) *T. brucei*. Transcripts for the stage-specific surface proteins procyclin, the variant or invariant surface glycoproteins VSG and ISG, the expression site-associated genes PAG and ESAG, and retrotransposon hotspot proteins (RHS) are colored as indicated. Dashed vertical lines: Fold changes (BS/PC) >4 or <-4. Dashed horizontal line: *p*-values <0.05.

Importantly, of the 705 lncRNAs included in the DGE analysis, 109 are differentially expressed (Figure 7a). Only four are up-regulated in procyclic-stage trypanosomes and one of them, the KS17gene_1079a, is flanked by the insect stage-specific procyclin genes Tb927.6.510 and Tb927.6.520. The remaining 105 lncRNAs are up-regulated in bloodstream parasites and 40% colocalize with VSG, ESAG or RHS genes or with pseudogenes. This suggests a co-regulation of nearly every 2nd lncRNA with the expression of stage-specific (surface) proteins making them prime candidates for further investigations. Finally, we experimentally confirmed the differential expression of 4 of the new lncRNAs (nt_4120.1, nt_256.1, nt_1759.1, nt_17.3) by qRT-PCR, which is shown in Figure 7b. The genomic context of the 4 lncRNAs as well as their sequencing coverage profiles for both bloodstream-stage and procyclic-stage trypanosomes is shown in Supplementary Figure S9.

**Figure 7.**
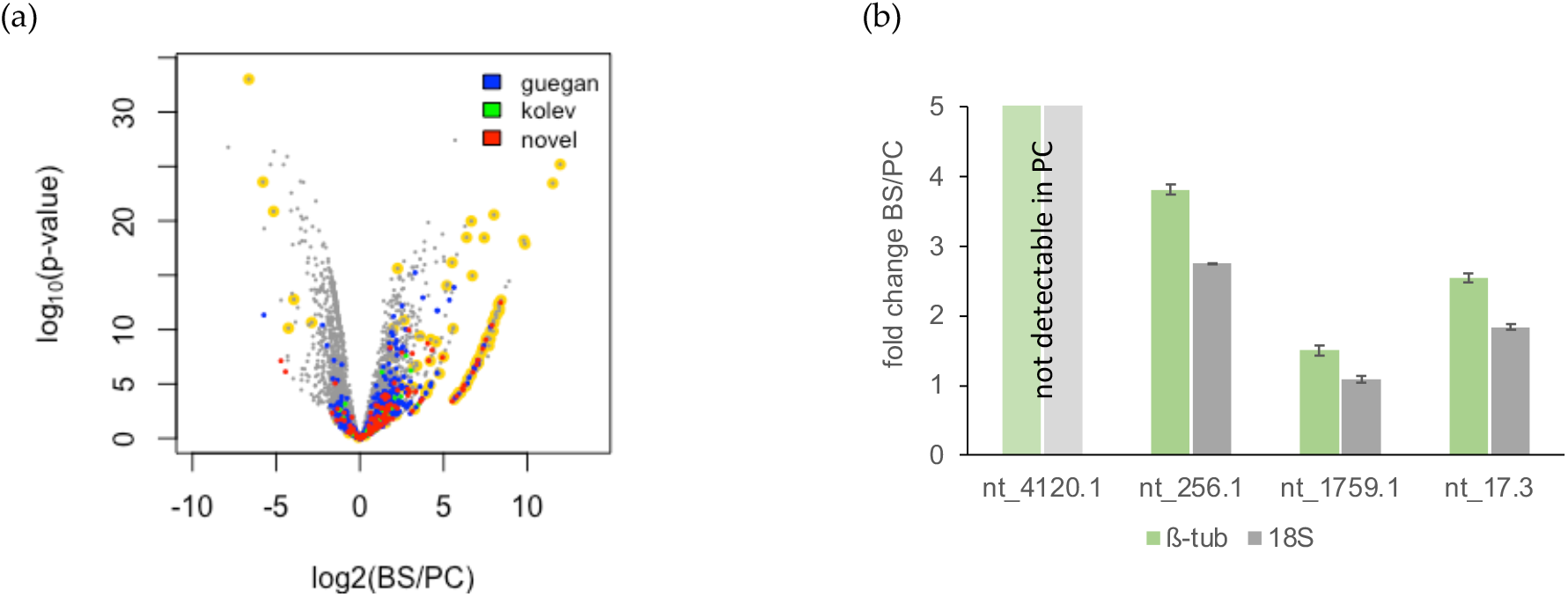
Differential expression of lncRNAs. (a) Volcano plot of differential gene expression between bloodstream-stage (BS) and procyclic-stage (PC) *T. brucei*. Novel lncRNA (this study) are in red. LncRNAs described in Guegan et al., 2022 and Kolev et al., 2010 are in blue and green. Transcripts for the stage-specific cell surface proteins procyclin, VSG, ISG and expression site-associated genes (ESAG) are highlighted in yellow. (b) Quantitative (q)RT-PCR results of transcript levels of the 4 novel lncRNAs: nt_4120.1, nt_256.1, nt_1759.1, and nt_17.3. Fold changes (BS/PC) were estimated by the ΔC_T_ method with either 18S rRNA (18S, grey) or β-tubulin (β-tub, green) as controls. Error bars are ±1SD. For nt_4120.1, no signal was detected in PC trypanosomes.

### 3.7 The mitochondrial transcriptome

Nearly the entire mitochondrial genome is covered by sequencing reads (Figure 8a). This is consistent with the observation that RNA polymerase occupies the entire maxicircle genome (Sement et al. 2018). In the coding region, gene boundaries are visible as discrete steps in the coverage profile, adjacent to poly(A) signature reads (Figure 8b). In the variable, *i.e., noncoding* region, the coverage pattern is rugged with no defined boundaries (Figure 8c). Transcripts from the noncoding region are only detected for the plus strand. However, based on the data, we cannot decide if the minus strand is not transcribed, or if only transcripts from the plus strand are captured because of its high A-nt content (Supplementary Figure S10). Di-cistronic reads are <0.1%, which is in agreement with monocistronic transcription (Sement et al. 2018). Steadystate transcript levels, estimated from the sequencing depth (Supplementary Figure S11), vary over 3 orders of magnitude and show the expected stage-specific signatures for edited RNAs (Figure 8b), which is further detailed below.

**Figure 8.**
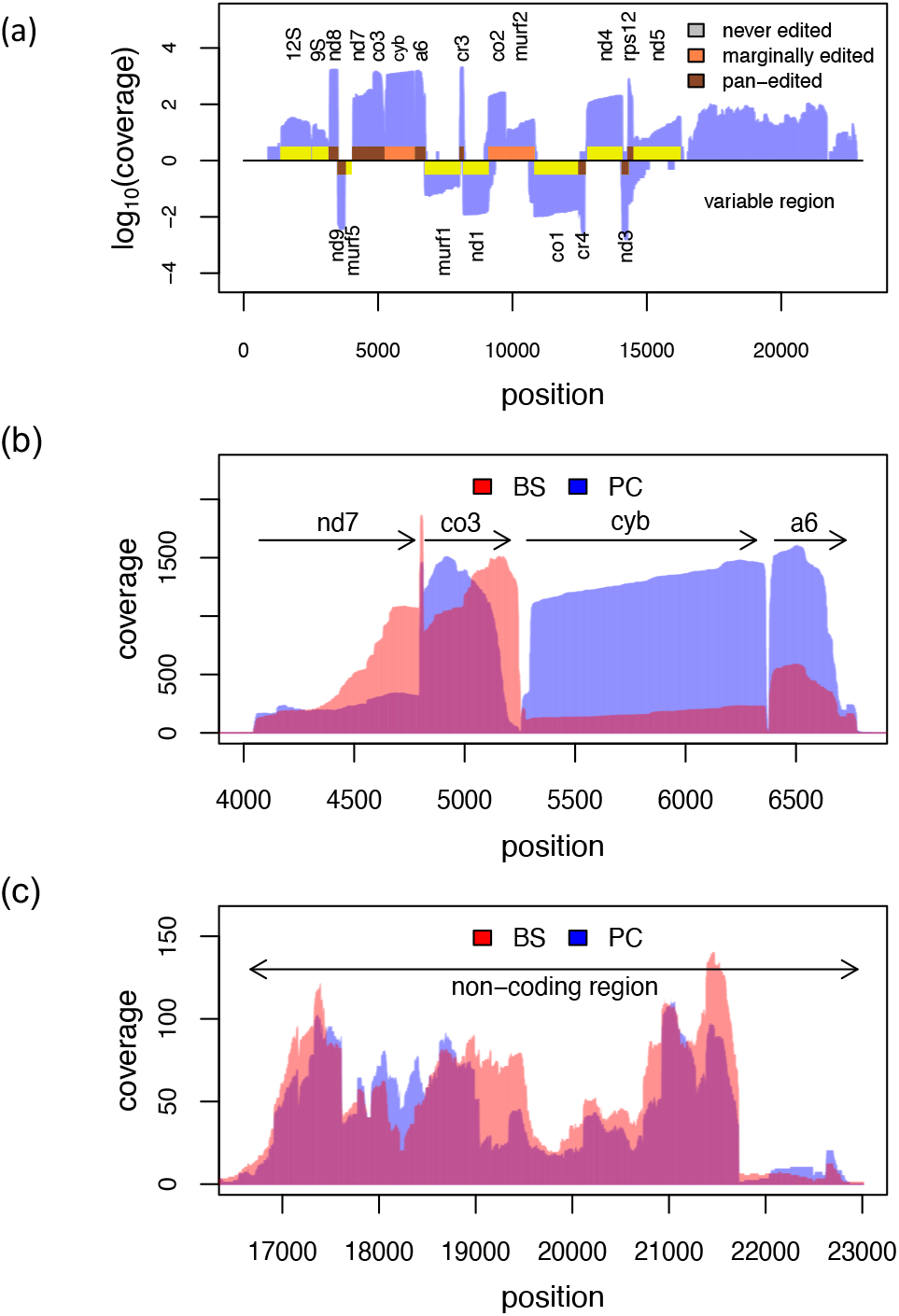
Mitochondrial gene expression. (a) Linear map of both strands of the *T. bruce*i mitochondrial (maxicircle) genome overlayed with the coverage profile of procyclic-stage trypanosomes (blue). Gene abbreviations are as in Aphasizheva et al., 2020. Never-edited genes are in yellow, marginally edited genes in orange and pan-edited genes in pink. (b) Overlay of the coverage profiles from bloodstream-stage (BS, red) and procyclic-stage (PC, blue) trypanosomes in the coding region from ND7 to A6. ND7=subunit 7 of the NADH dehydrogenase. CO3=cytochrome oxidase subunit 3. Cyb=cytochrome b. A6=subunit 6 of the mitochondrial ATPase. (c) Overlay of the coverage profiles from bloodstream-stage (BS, red) and procyclic-stage (PC, blue) trypanosomes in the non-coding (variable) region of the *T. brucei* maxicircle genome.

It should be noted that the absolute values are biased. For example, 9S and 12S ribosomal RNAs are expected to constitute the most abundant maxicircle transcripts. However, <1% of the reads map to the two rRNAs, possibly because they are 3’-oligouridylated (Adler et al., 1991) and therefore escape the poly(A)-enrichment step. Similarly, mitochondrial mRNAs have been shown to be 3’-AU tailed (Aphasizhev and Aphasizheva, 2011, Aphasizheva et al., 2020), which, as before, might have caused the depletion of certain transcripts during library preparation. This is likely also reflected in the relative fraction of polyadenylated mitochondrial reads. With a value of 55%, it is considerably lower compared to the fraction of poly(A)-tailed and *trans*-spliced RNAs (81%). The median length of the poly(A)-tails of all mitochondrial transcripts is about 30nt, with minor variations between different transcripts. The fraction of polyadenylation and the length of the poly(A)-tails are comparable in both life cycle stages (Supplementary Figure S12).

### 3.8 RNA editing

As shown in Figure 8b, the coverage profiles for never-edited or partially edited mitochondrial RNA also show the described 3’-end bias observed for nuclear-encoded transcripts (see Cyb mRNA in Figure 8b). However, highly edited, so-called pan-edited transcripts are characterized by an additional distortion towards the 3’-end, which is due to editing. Since the U deletion/U-insertion process proceeds from 3’ to 5’, only the 5’-regions of partially edited transcripts will align to the genomic sequences, which leads to a gradual drop-off in the DRS profiles towards the 3’-end (see CO3 and A6 transcripts in Figure 8b). Mapping all reads to the *T. brucei* mitochondrial transcriptome, including all fully edited sequences, was used to monitor the efficiency of the processing reaction (Figure 9). The fraction of reads representing fully edited mRNAs ranges from only 0.6% for the CO3 transcript to 25% for the RPS12 mRNA, which suggests that at steady state, fully edited mitochondrial transcripts are turned over rapidly. For genes encoding subunits of respiratory complex I (NADH dehydrogenase) no completely edited transcripts were detected in accordance with the findings of Opperdoes and Michels, 2008.

**Figure 9.**
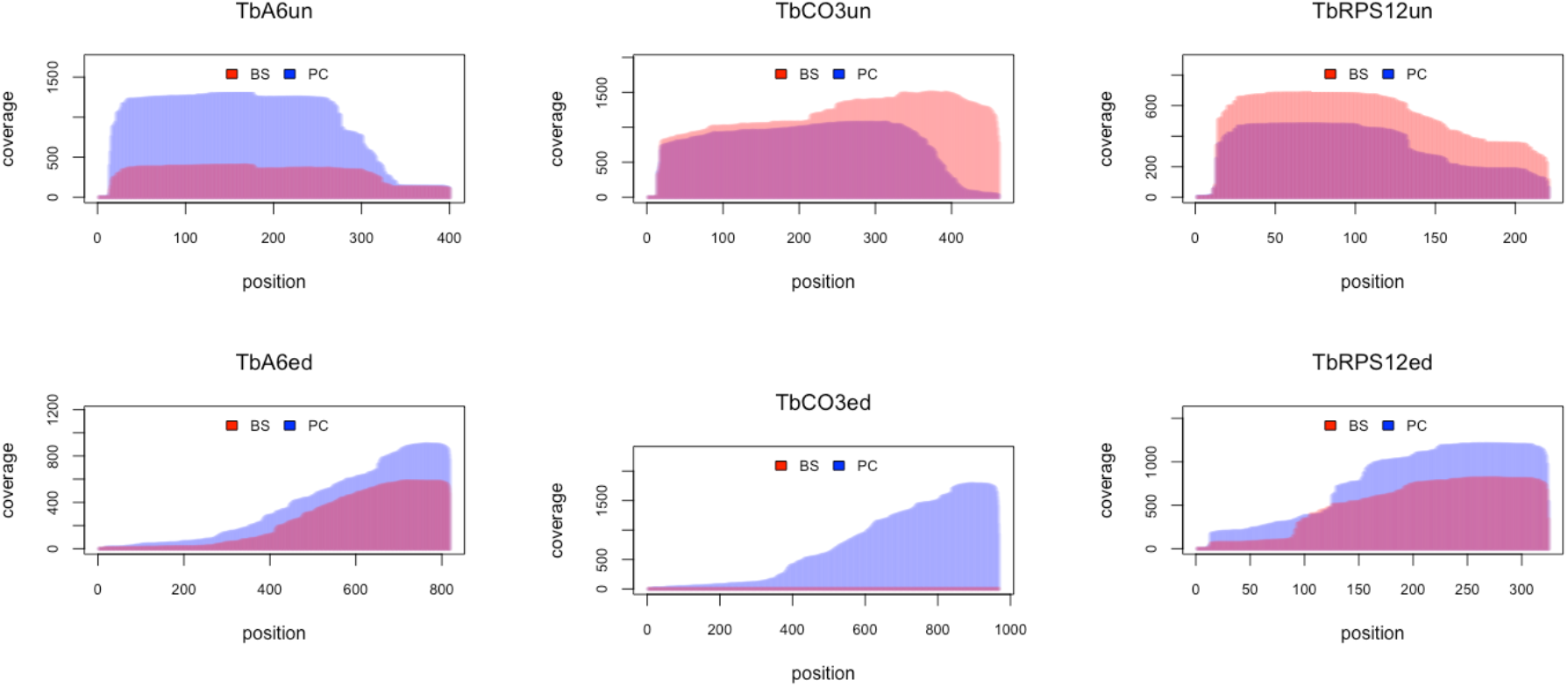
DRS captures the efficiency of mitochondrial RNA editing. Overlay profiles from bloodstream-stage (BS, red) and procyclic-stage (PC, blue) *T. brucei* (Tb) comparing the sequence coverage of unedited (un, top panel) and edited (ed, bottom panel) transcripts of the mitochondrial ATPase subunit 6 (A6), cytochrome oxidase 3 (CO3), and ribosomal protein S12 (RPS12).

## 5. Conclusions

In this work, we generated a set of genuine RNA sequencing data of the *T. brucei* transcriptome by using direct RNA sequencing on nanopore arrays. This was done to circumvent the reverse transcription and PCR amplification steps of short-read RNA sequencing to obtain strand-specific long-read RNA sequence information. We analyzed the two major stages of the parasite lifecycle using native poly(A)-tailed RNA and generated up to 0.7×10^6^ reads per library with a median read length of 1500nt and a maximal length >14500nt. A special emphasis was put on the inventory of lncRNAs as potential regulators of gene expression. Of the 1491 lncRNA genes annotated in the reference genome only 50% were confirmed by DRS. However, we identified 207 previously unknown lncRNAs, 109 of which are stage-specifically expressed with the majority (>96%) up-regulated in bloodstream-stage parasites. Strikingly, nearly every second of the newly identified lncRNA genes is localized proximal to a bloodstream-stage specific VSG, ESAG, or RHS gene, suggesting a co-regulation scenario perhaps by a similar mechanism. Furthermore, the DRS data provide insights into the complexity of the *T. brucei* transcriptome. This includes 1756 loci, which code for 5203 different transcript isoforms either representing splice variants of the same transcript or alternative or aberrant splicing products resulting in split or truncated exons or di- or poly-cistronic RNAs. And it includes a poly(A)-tail length plasticity between different RNA classes with a median length of 96nt for mRNAs, 117nt for lncRNAs, and only 29nt for mitochondrial mRNAs. Together, the data demonstrate that DRS can provide a more comprehensive and bias-free understanding of the *T. brucei* transcriptome. Provided that the current basecall accuracy and genome coverage limitations can be overcome (Jain et al., 2022), it should be possible to gain further insight into the plasticity of RNA processing reactions in *T. brucei*, which should accelerate our understanding of the RNA biology of the parasite.

## Supplementary Materials

Supplementary Figures S1-S12 and Supplementary Tables S1-S7 can be found in the online version of the article.

## Author Contributions

Conceptualization and funding acquisition H.U.G; experimentation, data analysis, and curation, E.K.; manuscript preparation E.K. and H.U.G. Both authors have read and agreed to the submitted version of the manuscript.

## Data Availability Statement

All sequencing data will be deposited in the European Nucleotide Archive (ENA) at EMBL-EBI.

## Acknowledgments

The authors thank Andreas Völker for growing and maintaining the *T. brucei* cell lines and Koraitim Malak and Maximillian Zick for their experimental help. Matthias Leeder is thanked for software installations and discussions.

## References

Adler, B.K.; Harris, M.E.; Bertrand, K.I.; Hajduk, S.L. Modification of *Trypanosoma brucei* mitochondrial rRNA by posttranscriptional 3’ polyuridine tail formation. Mol. Cell. Biol. 1991, 11, 5878–5884.

Aird, D.; Ross, M.G.; Chen, W.S.; Danielsson, M.; Fennell, T.; Russ, C.; Jaffe, D.B.; Nusbaum, C.; Gnirke, A. Analyzing and minimizing PCR amplification bias in Illumina sequencing libraries. Genome Biol. 2011, 12, R18.

Aphasizhev, R.; Aphasizheva, I. Mitochondrial RNA processing in trypanosomes. Res. Microbiol. 2011, 162, 655–663.

Aphasizheva, I.; Alfonzo, J.; Carnes, J.; Cestari, I.; Cruz-Reyes, J.; Göringer, H.U.; Hajduk, S.; Lukeš, J.; Madison-Antenucci, S.; Maslov, D.A.; McDermott, S.M.; Ochsenreiter, T.; Read, L.K.; Salavati, R.; Schnaufer, A.; Schneider, A.; Simpson, L.; Stuart, K.; Yurchenko, V.; Zhou, Z.H.; Zíková, A.; Zhang, L.; Zimmer, S.; Aphasizhev, R. Lexis and Grammar of Mitochondrial RNA Processing in Trypanosomes. Trends Parasitol. 2020, 36, 337–355.

Bringaud, F.; Biteau, N.; Melville, S.E.; Hez, S.; El-Sayed, N.M.; Leech, V.; Berriman, M.; Hall, N.; Donelson, J.E.; Baltz, T. A new, expressed multigene family containing a hot spot for insertion of retroelements is associated with polymorphic subtelomeric regions of *Trypanosoma brucei. Eukaryot*. Cell. 2002, 1, 137–151. Erratum in: *Eukaryot. Cell* 2002, 1, 305.

Bringaud, F.; Biteau, N.; Zuiderwijk, E.; Berriman, M.; El-Sayed, N.M.; Ghedin, E.; Melville, S.E.; Hall, N.; Baltz, T. The ingi and RIME non-LTR retrotransposons are not randomly distributed in the genome of *Trypanosoma brucei*. Mol. Biol. Evol. 2004, 21, 520–528.

Brun, R.; Schönenberger, M. Cultivation and *in vitro* cloning or procyclic culture forms of *Trypanosoma brucei* in a semi-defined medium. Acta Trop. 1979, 36, 289–292.

Byrne, A.; Beaudin, A.E.; Olsen, H.E.; Jain, M.; Cole, C.; Palmer, T.; DuBois, R.M.; Forsberg, E.C.; Akeson, M.; Vollmers, C. Nanopore long-read RNAseq reveals widespread transcriptional variation among the surface receptors of individual B cells. Nat. Commun. 2017, 8, 16027.

Chomczynski, P.; Sacchi, N. Single-step method of RNA isolation by acid guanidinium thiocyanate-phenol-chloroform extraction. Anal. Biochem. 1987, 162, 156–159.

Christiano, R.; Kolev, N.G.; Shi, H.; Ullu, E.; Walther, T.C.; Tschudi, C. The proteome and transcriptome of the infectious metacyclic form of *Trypanosoma brucei* define quiescent cells primed for mammalian invasion. Mol. Microbiol. 2017, 106, 74–92.

Clayton, C. Regulation of gene expression in trypanosomatids: living with polycistronic transcription. Open Biol. 2019, 9, 190072.

Clayton, C.; Shapira, M. Post-transcriptional regulation of gene expression in trypanosomes and leishmanias. Mol Biochem Parasitol. 2007, 156, 93–101.

Cross, G.A. Identification, purification and properties of clone-specific glycoprotein antigens constituting the surface coat of *Trypanosoma brucei*. Parasitology 1975, 71, 393–417.

De Rycker, M.; Wyllie, S.; Horn, D.; Read, K.D.; Gilbert, I.H. Anti-trypanosomatid drug discovery: progress and challenges. Nat. Rev. Microbiol. 2023, 21, 35–50.

Dawe, H.R.; Shaw, M.K.; Farr, H.; Gull, K. The hydrocephalus inducing gene product, Hydin, positions axonemal central pair microtubules. BMC Biology 2007, 5, 33.

Fickett, J.W.; Tung, C.S. Assessment of protein coding measures. Nucleic Acids Res. 1992, 20, 6441–6450.

Fort, R.S.; Chavez, S.; Trinidad Barnech, J.M.; Oliveira-Rizzo, C.; Smircich, P.; Sotelo-Silveira, J.R.; Duhagon, M.A. Current Status of Regulatory Non-Coding RNAs Research in the Tritryp. Noncoding RNA 2022, 8, 54.

Garalde, D.R.; Snell, E.A.; Jachimowicz, D.; Sipos, B.; Lloyd, J.H.; Bruce, M.; Pantic, N.; Admassu, T.; James, P.; Warland, A.; Jordan, M.; Ciccone, J.; Serra, S.; Keenan, J.; Martin, S.; McNeill, L.; Wallace, E.J.; Jayasinghe, L.; Wright, C.; Blasco, J.; Young, S.; Brocklebank, D.; Juul, S.; Clarke, J.; Heron, A.J.; Turner, D.J. Highly parallel direct RNA sequencing on an array of nanopores. Nat. Methods 2018, 15, 201–206.

Gleeson, J.; Leger, A.; Prawer, Y.D.J.; Lane, T.A.; Harrison, P.J.; Haerty, W.; Clark, M.B. Accurate expression quantification from nanopore direct RNA sequencing with NanoCount. Nucleic Acids Res. 2022, 50, e19.

Guegan, F.; Rajan, K.S.; Bento, F.; Pinto-Neves, D.; Sequeira, M.; Gumińska, N.; Mroczek, S.; Dziembowski, A.; Cohen-Chalamish, S.; Doniger, T.; Galili, B.; Estévez, A.M.; Notredame, C.; Michaeli, S.; Figueiredo, L.M. A long noncoding RNA promotes parasite differentiation in African trypanosomes. Sci. Adv. 2022, 8, eabn2706.

Han, S.; Liang, Y.; Ma, Q.; Xu, Y.; Zhang, Y.; Du, W.; Wang, C.; Li, Y. LncFinder: an integrated platform for long noncoding RNA identification utilizing sequence intrinsic composition, structural information and physicochemical property. Brief Bioinform. 2019, 20, 2009–2027.

Hirumi, H.; Hirumi, K. Continuous cultivation of *Trypanosoma brucei* blood stream forms in a medium containing a low concentration of serum protein without feeder cell layers. J. Parasitol. 1989, 75, 985–989.

Houseley, J.; Tollervey, D. Apparent non-canonical *trans*-splicing is generated by reverse transcriptase *in vitro*. PLoS One 2010, 5, e12271.

Jan, C.H.; Friedman, R.C.; Ruby, J.G.; Bartel, D.P. Formation, regulation and evolution of *Caenorhabditis elegans* 3’UTRs. Nature 2011, 469, 97–101.

Jain, M.; Abu-Shumays, R.; Olsen, H.E.; Akeson, M. Advances in nanopore direct RNA sequencing. Nat. Methods 2022, 19, 1160–1164.

Jenjaroenpun, P.; Wongsurawat, T.; Pereira, R.; Patumcharoenpol, P.; Ussery, D.W.; Nielsen, J.; Nookaew, I. Complete genomic and transcriptional landscape analysis using third-generation sequencing: a case study of *Saccharomyces cerevisiae* CEN.PK113-7D. Nucleic Acids Res. 2018, 46, e38.

Kang, Y.J.; Yang, D.C.; Kong, L.; Hou, M.; Meng, Y.Q.; Wei, L.; Gao, G. CPC2: a fast and accurate coding potential calculator based on sequence intrinsic features. Nucleic Acids Res. 2017, 45, W12–W16.

Kaufer, A.; Stark, D.; Ellis, J. A review of the systematics, species identification and diagnostics of the *Trypanosomatidae* using the maxicircle kinetoplast DNA: from past to present. Int. J. Parasitol. 2020, 50, 449–460.

Kolev, N.G.; Franklin, J.B.; Carmi, S.; Shi, H.; Michaeli, S.; Tschudi, C. The transcriptome of the human pathogen *Trypanosoma brucei* at single-nucleotide resolution. PLoS Pathog. 2010, 6, e1001090.

Kolev, N.G.; Rajan, K.S.; Tycowski, K.T.; Toh, J.Y.; Shi, H.; Lei, Y.; Michaeli, S.; Tschudi, C. The vault RNA of *Trypanosoma brucei* plays a role in the production of *trans*-spliced mRNA. J. Biol. Chem. 2019, 294, 15559–15574.

Kolev, N.G.; Ullu, E.; Tschudi, C. The emerging role of RNA-binding proteins in the life cycle of *Trypanosoma brucei*. Cell. Microbiol. 2014, 16, 482–489.

Kong, L.; Zhang, Y.; Ye, Z.Q.; Liu, X.Q.; Zhao, S.Q.; Wei, L.; Gao, G. CPC: assess the protein-coding potential of transcripts using sequence features and support vector machine. Nucleic Acids Res. 2007, 35: W345–W349.

Koslowsky, D.; Sun, Y.; Hindenach, J.; Theisen, T.; Lucas, J. The insect-phase gRNA transcriptome in *Trypanosoma brucei*. Nucleic Acids Res. 2014, 42, 1873–1886.

Krause, M.; Niazi, A.M.; Labun, K.; Torres Cleuren, Y.N.; Müller, F.S.; Valen, E. tailfindr: alignment-free poly(A) length measurement for Oxford Nanopore RNA and DNA sequencing. RNA 2019, 25, 1229–1241.

Li, H.; Durbin, R. Fast and accurate short read alignment with Burrows-Wheeler transform. Bioinformatics 2009, 25, 1754–1760.

Li, H. Minimap2: pairwise alignment for nucleotide sequences. Bioinformatics 2018, 34, 3094–3100.

Liao, Y.; Smyth, G.K.; Shi, W. The R package Rsubread is easier, faster, cheaper and better for alignment and quantification of RNA sequencing reads. Nucleic Acids Res. 2019, 47, e47.

Liao, Y.; Smyth, G.K.; Shi, W. featureCounts: an efficient general purpose program for assigning sequence reads to genomic features. Bioinformatics 2014, 30, 923–930.

Livak, K.J.; Schmittgen, T.D. Analysis of relative gene expression data using real-time quantitative PCR and the 2(- Delta Delta C(T)) Method. Methods 2001, 25, 402–408.

Loman, N.J.; Quick, J.; Simpson, J.T. A complete bacterial genome assembled de novo using only nanopore sequencing data. Nat. Methods 2015, 12, 733–735.

Lorenz, R.; Bernhart, S.H.; Höner zu Siederdissen, C.; Tafer, H.; Flamm, C.; Stadler, P.F.; Hofacker, I.L. ViennaRNA Package 2.0. Algorithms Mol. Biol. 2011, 6, 1–26.

Martínez-Calvillo, S.; Florencio-Martínez, L.E.; Nepomuceno-Mejía, T. Nucleolar Structure and Function in Trypanosomatid Protozoa. Cells 2019, 8, 421.

Michaeli, S. *Trans*-splicing in trypanosomes: machinery and its impact on the parasite transcriptome. Future Microbiol. 2011, 6, 459–474.

Michaeli, S.; Podell, D.; Agabian, N.; Ullu, E. The 7SL RNA homologue of *Trypanosoma brucei* is closely related to mammalian 7SL RNA. Mol. Biochem. Parasitol. 1992, 51, 55–64.

Maier, K.C.; Gressel, S.; Cramer, P.; Schwalb, B. Native molecule sequencing by nano-ID reveals synthesis and stability of RNA isoforms. Genome Res. 2020, 30, 1332–1344.

Minshall, N.; Git, A. Enzyme- and gene-specific biases in reverse transcription of RNA raise concerns for evaluating gene expressio. Sci. Rep. 2020, 10, 8151.

Mourão, K.; Schurch, N.J.; Lucoszek, R.; Froussios, K.; MacKinnon, K.; Duc, C.; Simpson, G.; Barton, G.J. Detection and mitigation of spurious antisense expression with RoSA. F1000Research 2019, 8, 819

Opperdoes, F.R.; Michels, P.A. Complex I of Trypanosomatidae: does it exist? Trends Parasitol. 2008, 24, 310–317.

Parekh, S.; Ziegenhain, C.; Vieth, B.; Enard, W.; Hellmann, I. The impact of amplification on differential expression analyses by RNA-seq. Sci. Rep. 2016, 6, 25533.

Quinlan, A.R. BEDTools: The Swiss-Army Tool for Genome Feature Analysis. Curr. Protoc. Bioinformatics 2014, 47, 11.12.1-34.

Quinlan, A.R.; Hall, I.M. BEDTools: a flexible suite of utilities for comparing genomic features. Bioinformatics 2010, 26, 841–842.

Rajan, K.S.; Doniger, T.; Cohen-Chalamish, S.; Rengaraj, P.; Galili, B.; Aryal, S.; Unger, R.; Tschudi, C.; Michaeli, S. Developmentally Regulated Novel Non-coding Anti-sense Regulators of mRNA Translation in *Trypanosoma brucei*. iScience 2020, 23, 101780.

Robinson, M.D.; McCarthy, D.J.; Smyth, G.K. edgeR: a Bioconductor package for differential expression analysis of digital gene expression data. Bioinformatics. 2010, 26, 139–140.

Sandhu, R.; Sanford, S.; Basu, S.; Park, M.; Pandya, U.M.; Li, B.; Chakrabarti, K. A *trans*-spliced telomerase RNA dictates telomere synthesis in *Trypanosoma brucei*. Cell Res. 2013, 23, 537–551.

Sement, F.M.; Suematsu, T.; Zhang, L.; Yu, T.; Huang, L.; Aphasizheva, I.; Aphasizhev, R. Transcription initiation defines kinetoplast RNA boundaries. Proc. Natl. Acad. Sci. USA 2018, 115, e10323–E10332.

Siegel, T.N.; Hekstra, D.R.; Wang, X.; Dewell, S.; Cross, G.A. Genome-wide analysis of mRNA abundance in two life-cycle stages of *Trypanosoma brucei* and identification of splicing and polyadenylation sites. Nucleic Acids Res. 2010, 38, 4946–4957.

Steijger, T.; Abril J.F.; Engström, P.G.; Kokocinski, F.; RGASP Consortium; Hubbard, T.J.; Guigó, R.; Harrow, J.; Bertone, P. Assessment of transcript reconstruction methods for RNA-seq. Nat. Methods 2013, 10, 1177–1184.

Statello, L.; Guo, C.J.; Chen, L.L.; Huarte, M. Gene regulation by long non-coding RNAs and its biological functions. Nat. Rev. Mol. Cell. Biol. 2021, 22, 96–118.

Veitch, N.J.; Johnson, P.C.; Trivedi, U.; Terry, S.; Wildridge, D.; MacLeod, A. Digital gene expression analysis of two life cycle stages of the human-infective parasite, *Trypanosoma brucei gambiense* reveals differentially expressed clusters of co-regulated genes. BMC Genomics. 2010, 11, 124.

Wick, R.R.; Judd, L.M.; Holt, K.E. Performance of neural network basecalling tools for Oxford Nanopore sequencing. Genome Biology 2019, 20, 129.

Workman, R.E.; Tang, A.D.; Tang, P.S.; Jain, M.; Tyson, J.R.; Razaghi, R.; Zuzarte, P.C.; Gilpatrick, T.; Payne, A.; Quick, J.; Sadowski, N.; Holmes, N.; de Jesus, J.G.; Jones, K.L.; Soulette, C.M.; Snutch, T.P.; Loman, N.; Paten, B.; Loose, M.; Simpson, J.T.; Olsen, H.E.; Brooks, A.N.; Akeson, M.; Timp, W. Nanopore native RNA sequencing of a human poly(A) transcriptome. Nat Methods 2019, 16, 1297–1305. Erratum in: *Nat Methods* 2020, 7, 114.

