## Supplementary Material for "Nanopore-based direct RNA sequencing of the *Trypanosoma brucei* transcriptome identifies novel lncRNAs"

**Table of Contents**

**1. Supplementary Figures S1 to S12**
**2. Supplementary Tables S1 to S7**

### Supplementary Figures

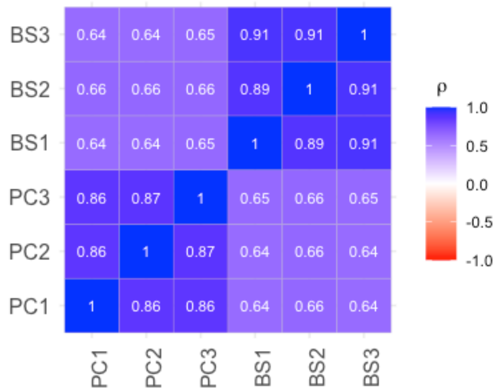

**Supplementary Figure S1**. Pairwise comparison of gene expression levels of all DRS sequencing libraries. BS1, BS2, BS3: libraries from bloodstream-stage *T. brucei*. PC1, PC2, PC3: libraries from procyclic-stage parasites. Gene expression levels were determined using featureCounts from the R-package Rsubread (Liao et al., 2014; Liao et al., 2019). *p*=Spearman rank correlation coefficient. Exons with low expression levels (<6 reads) were not considered.

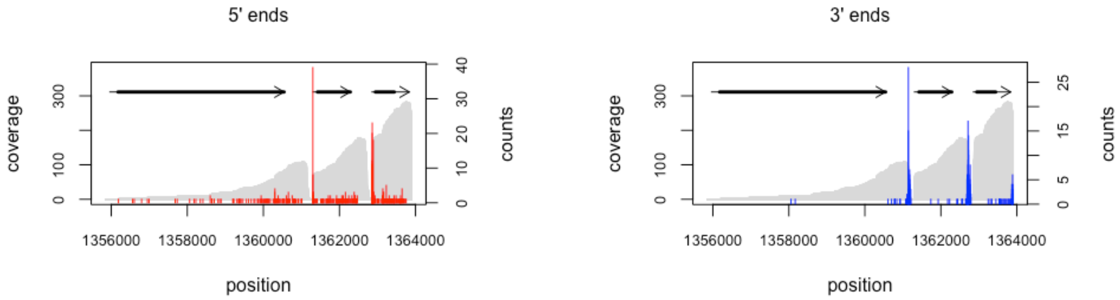

**Supplementary Figure S2.** Distribution of transcript start- and end-positions. Coverage profiles (grey) from bloodstream-stage trypanosomes for the genomic region Tb927_07_v5.1:1356000-13764000. Transcript start sites are in red and transcript end-positions are in blue. Grey arrows are annotated exons. Bold, black lines are coding sequences.

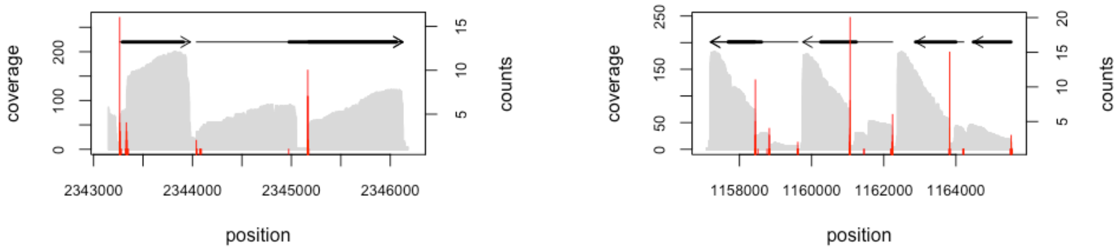

**Supplementary Figure S3**. Examples of splice acceptor (SAS) sites located within annotated coding sequences. Coverage profiles (grey) of two representative regions in the *T. brucei* genome (Tb927_09_v5.1:2343000-2346000(+), Tb927_08_v5.1:1157080-1165560(-), derived from bloodstream-stage trypanosomes. SAS sites, identified by the presence of a 5'-spliced leader (SL)-sequence are shown in red. Peak heights indicate read numbers. Grey arrows are annotated exons. Bold black lines are coding sequences.

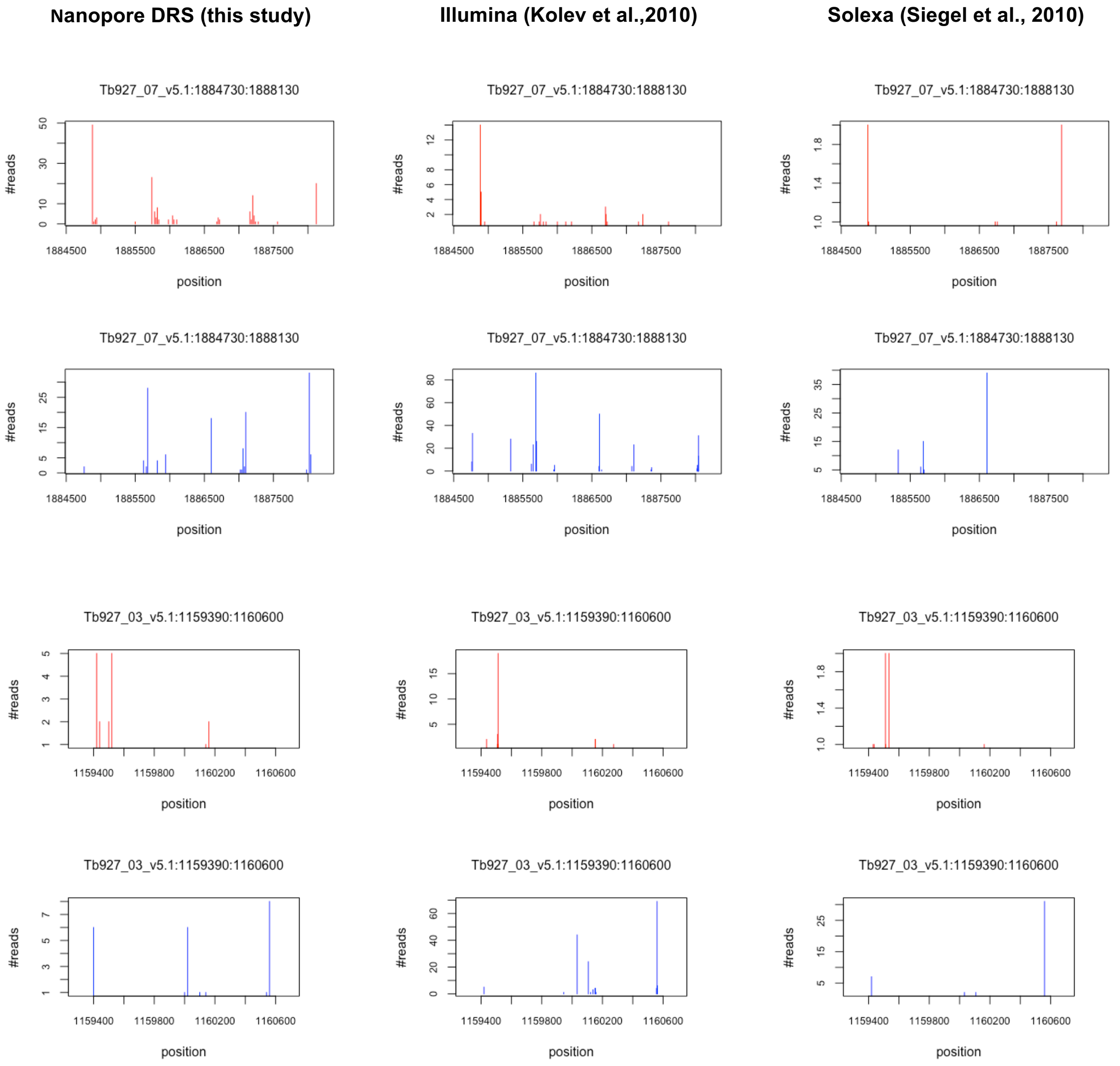

**Supplementary Figure S4.** Side-by-side comparison of mapped splice acceptor (SAS, red) and polyadenylation (PAS, blue) sites for two selected regions of the *T. brucei* genome, derived from different sequencing methods. Left: Nanopore-based DRS (this study). Center: Short-read Illumina sequencing (Kolev et al., 2010). Right: Short-read Solexa sequencing (Siegel et al., 2010). For the DRS data, the number of reads (start- or end-positions) was summed up over a window of 20nt.

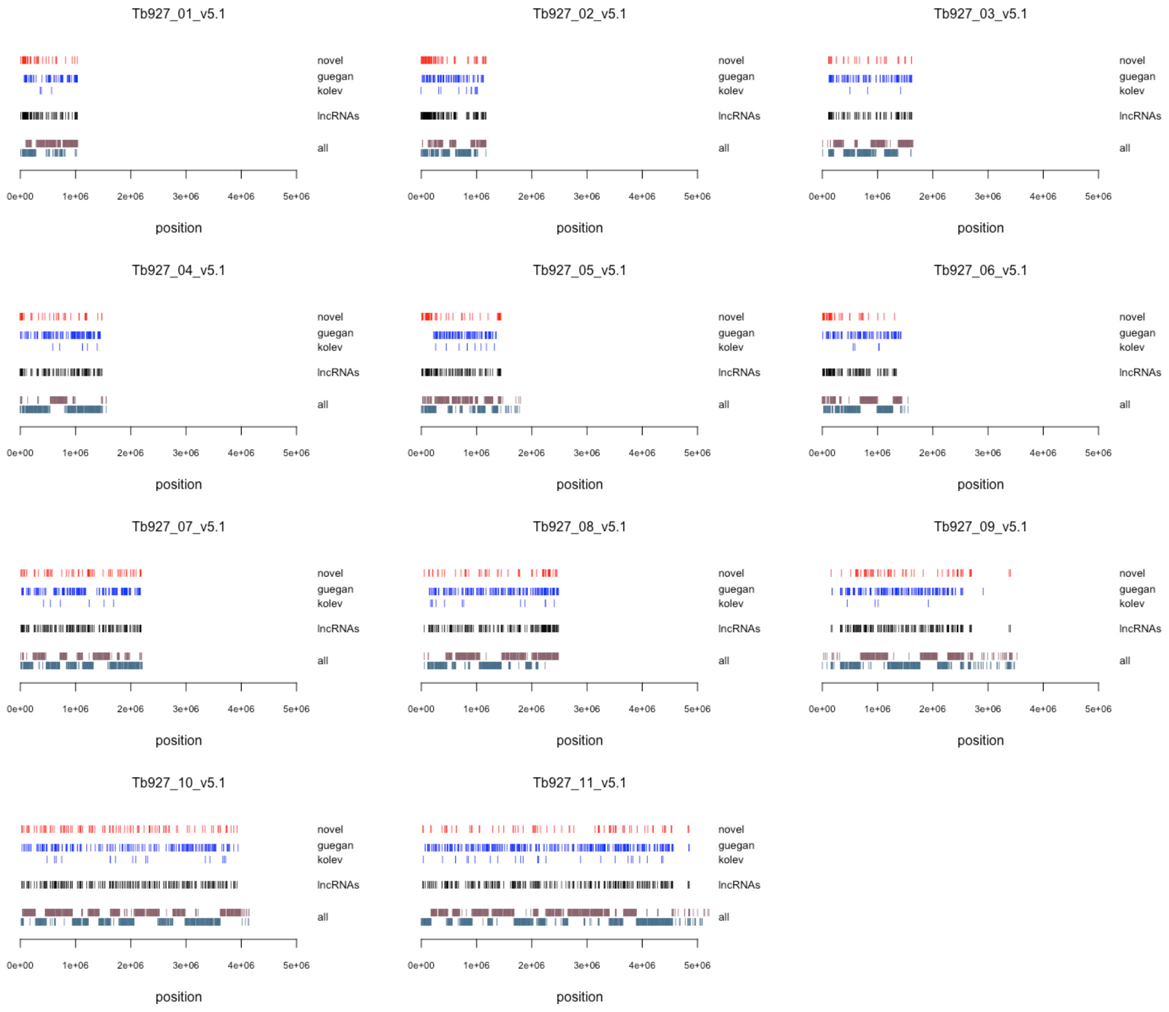

**Supplementary Figure S5.** Localization of all newly identified lncRNA genes (this study, red) on the 11 Mbp-size chromosomes of *T. brucei* (top to bottom) in comparison to the published sequences by Guegan et al., 2022 (blue) and Kolev et al., 2010 (grey). Black: all three datasets combined. All: strand-specific representation of all lncRNA genes. Top: plus strand. Bottom: minus strand.

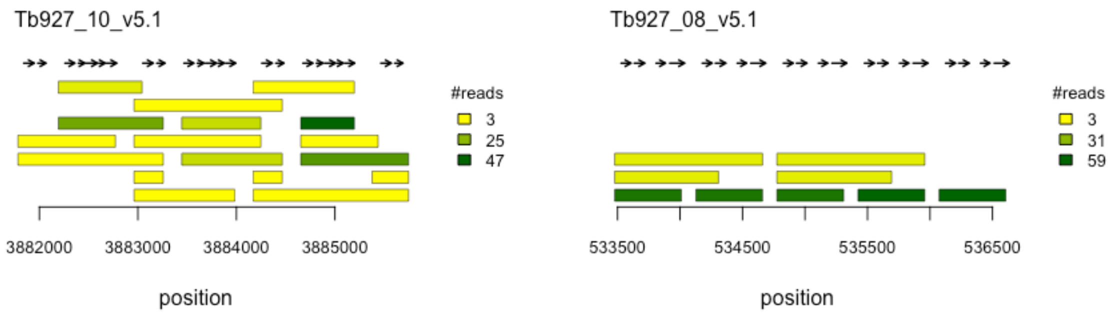

**Supplementary Figure S6.** Non-coding RNAs as precursors of snoRNAs. Examples from two regions of the *T. brucei* genome on chromosome 10 (left) and chromosome 8 (right). Horizontal bars are lncRNAs. Colors indicate the number of reads as indicated. Annotated snoRNAs are shown as black arrows.

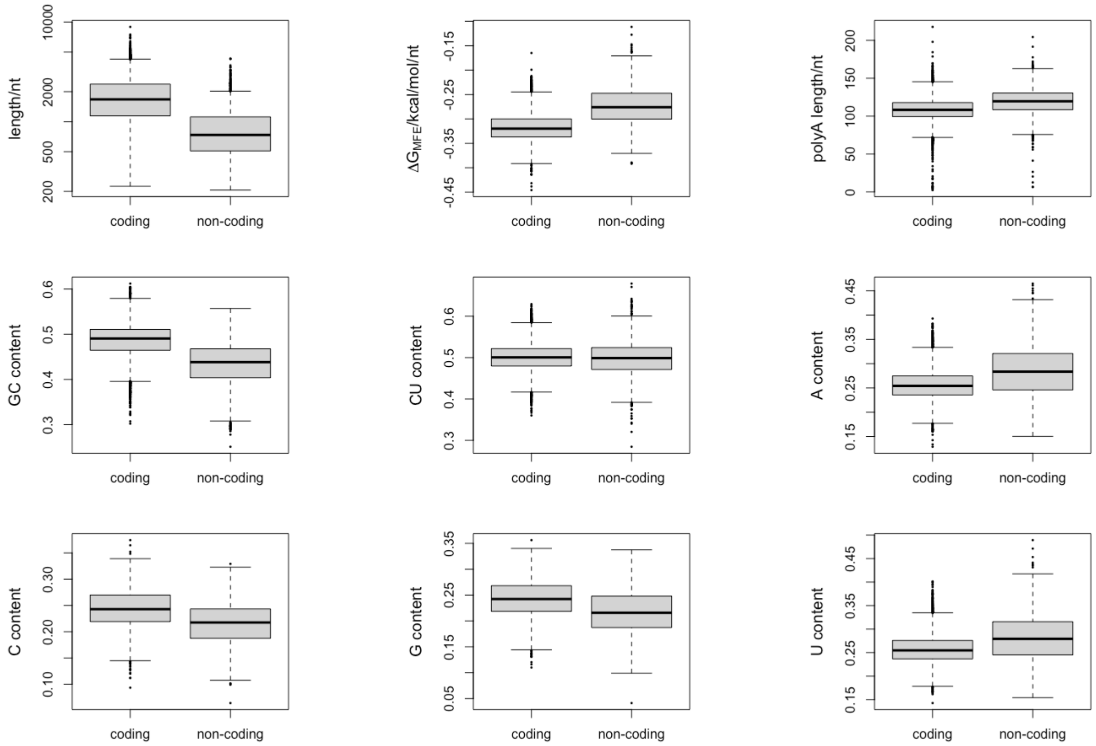

**Supplementary Figure S7**. Box-plot comparison of coding and noncoding transcripts from *T. brucei* for sequence and 2D-structure-specific parameters including nucleotide (nt) length, nt-content (A, C, G, U), GC- and CU-content, poly(A)-tail length, and thermodynamic stability (ΔG) of the minimal free energy structure (MFE). Transcripts were identified based on full-length reads as detailed in the Materials and Methods section. The coding potential was assessed by CPC2 (Kang et al., 2017) and LncFinder (Han et al., 2019). Only transcripts predicted as noncoding by both software tools were considered. ΔG-calculations were performed using RNAfold (Lorenz et al., 2011). Poly(A)-tails were assessed with the help of nanopolish (Workman et al., 2019).

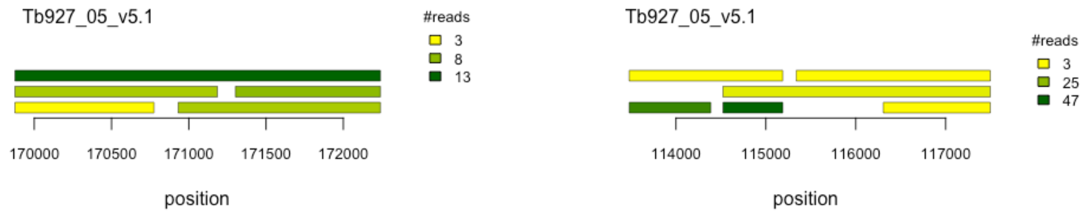

**Supplementary Figure S8.** Complex clustering of lncRNA genes. Examples from two different regions on chromosome 5 of the *T. brucei* genome are shown. Horizontal bars represent lncRNAs with colors indicating the number of sequencing reads.

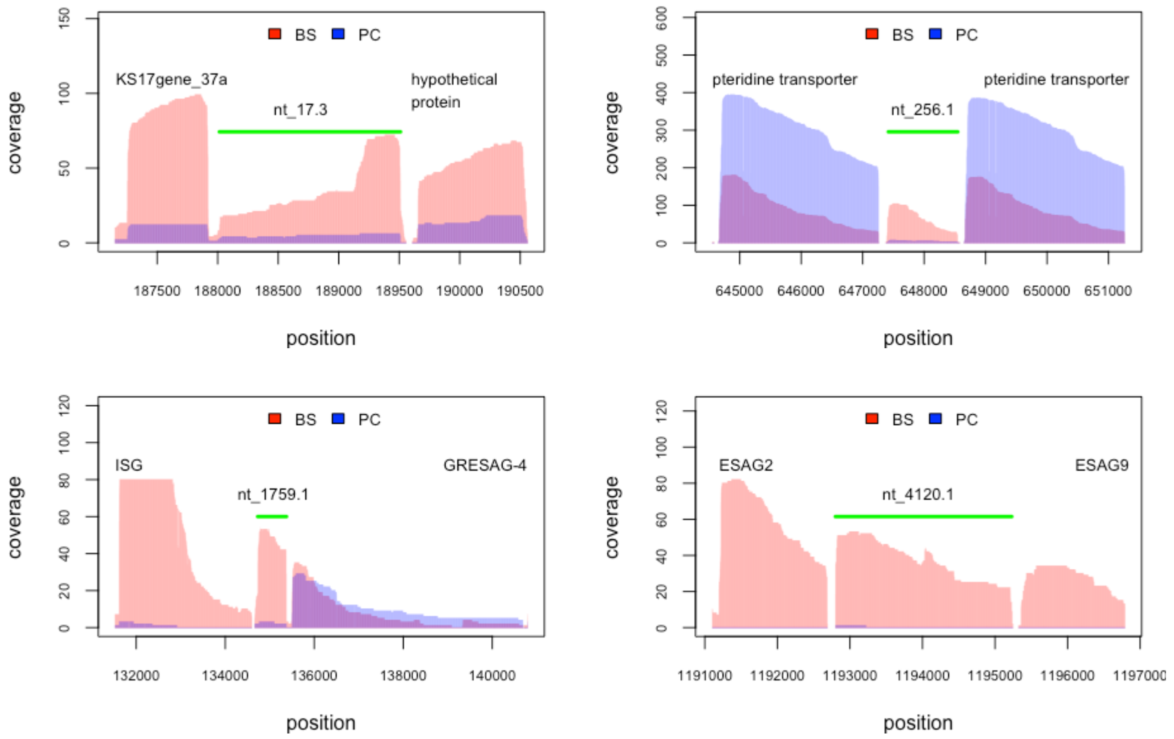

**Supplementary Figure S9.** Genomic context and differential expression of 4 newly identified lncRNAs (nt_17.3, nt_256.1, nt_1759.1, nt_4120.1) shown as green lines. Overlay plots of sequencing coverage profiles from bloodstream-stage (BS, red) and procyclic-stage (PC, blue) trypanosomes are shown. Flanking genes include the lncRNA gene K1717gene_37a, pteridine transporter genes, an invariant surface glycoprotein (ISG), the expression site-associated genes (ESAG) 2 and 9 and the glycine-rich expression site-associated gene 4 (GRESAG-4).

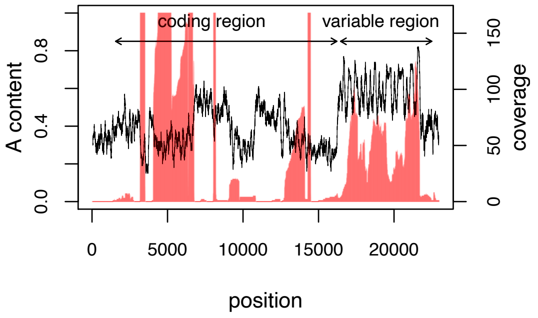

**Supplementary Figure S10.** Transcript coverage in the variable (noncoding) region of the *T. brucei* mitochondrial genome correlates with the local A-nucleotide content. Overlay plot of a DRS sequencing coverage profile (red) with the local A-nt content (black) of the plus strand of the mitochondrial genome. The data are derived from all procyclic-stage DRS libraries and the A-nt content was calculated over window of 100nt.

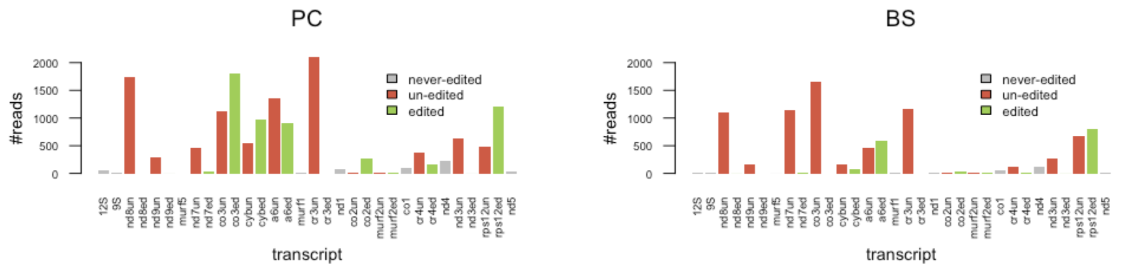

**Figure S11.** Steady state levels of mitochondrial transcripts in procyclic-stage (PC) and bloodstream-stage (BS) *T. brucei*. Reads were mapped to the mitochondrial transcriptome and steady state RNA levels were estimated from the number of sequencing reads. Grey=never-edited transcripts. Red=un-edited RNAs. Green=edited RNAs. Gene annotations are as in Aphasizheva et al., 2020.

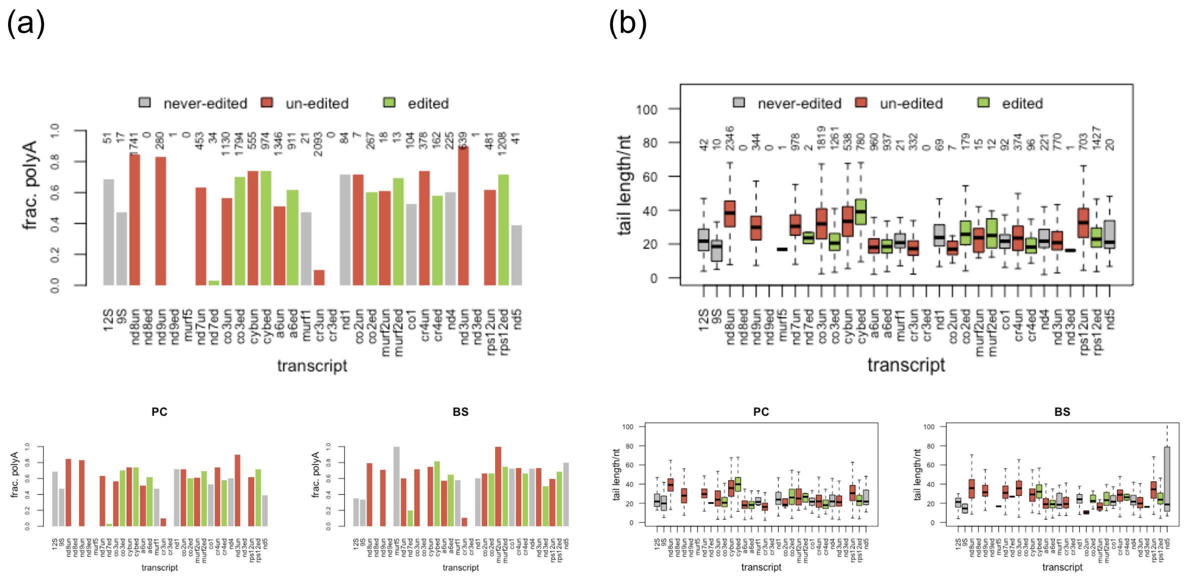

**Supplementary Figure S12**. Polyadenylation of mitochondrial RNAs. (a) The fraction (frac.) of sequencing reads holding a poly(A )-tail is plotted for all *T. brucei* mitochondrial transcripts. Transcript abbreviations are as in Aphasizheva et al., 2020. The total number of reads is indicated by the numbers above each bar. (b) Box-plot of the poly(A)-tail length of all mitochondrial transcripts. Upper panels: Data derived from the combined reads all DRS libraries (bloodstream-stage (BS) plus procyclic-stage (PC). Lower panels: The same data but separated for PC and BS trypanosomes. Grey=never-edited transcripts. Red=un-edited transcripts. Green=edited transcripts.

#### Supplementary Tables

**Supplementary Table S1.** Read statistics of the DRS sequencing data. Base calling was performed post-sequencing using the high accuracy (HAC) calling model of Guppy (Wick et al., 2019). Only reads with a mean per base quality score (*q*-score) >7 were used for further analysis.

| | **procyclic-stage *T. brucei*** | | | **bloodstream-stage *T. brucei*** | | |
|---|---|---|---|---|---|---|
| | **rep1** | **rep2** | **rep3** | **rep1** | **rep2** | **rep3** |
| **reads total** | 2.85E+05 | 3.37E+05 | 3.71E+05 | 6.02E+05 | 6.66E+05 | 4.77E+05 |
| **mean qscore >=7** | 2.64E+05 | 3.10E+05 | 3.43E+05 | 5.39E+05 | 5.67E+05 | 4.43E+05 |
| | 92% | 92% | 92% | 89% | 85% | 93% |
| **median qscore** | | | | | | |
| pass | 10.43 | 9.86 | 10.32 | 10.25 | 9.65 | 10.46 |
| fail | 5.32 | 5.95 | 5.80 | 5.78 | 5.50 | 5.62 |
| **median length** | | | | | | |
| pass | 779 | 811 | 786 | 825 | 666 | 750 |
| fail | 283 | 451 | 378 | 319 | 198 | 244 |
| **max length** | | | | | | |
| pass | 11784 | 12690 | 14262 | 12971 | 8697 | 7912 |
| fail | 14139 | 10395 | 25252 | 21789 | 18425 | 15827 |

**Supplementary Table S2.** Transcript mapping statistics. Reads with *q*-scores >7 were aligned to a composite reference genome (complete rerefence) consisting of the annotated chromosomes of *T. brucei* TREU927 (TryTrypDBv52), the *T. brucei* mitochondrial (maxicircle) genome (NCBI: nucleotide database, accNo M94286.1), and the sequence of yeast ENO2 used as reference calibration sequence (RCS) during library preparation.

| | **procyclic-stage *T. brucei*** | | | **bloodstream-stage *T. brucei*** | | |
|---|---|---|---|---|---|---|
| | **rep1** | **rep2** | **rep3** | **rep1** | **rep2** | **rep3** |
| input | 2.64E+05 | 3.10E+05 | 3.43E+05 | 5.39E+05 | 5.67E+05 | 4.43E+05 |
| **# alignments** | | | | | | |
| all alignments | 4.18E+05 | 4.93E+05 | 5.38E+05 | 8.17E+05 | 8.60E+05 | 6.80E+05 |
| unique reads | 2.59E+05 | 3.03E+05 | 3.35E+05 | 5.04E+05 | 5.24E+05 | 4.16E+05 |
| **RCS** | | | | | | |
| unique reads | 9901 | 12020 | 9803 | 16319 | 4802 | 19147 |
| **maxicircle genome** | | | | | | |
| unique reads | 3641 | 3385 | 4440 | 2522 | 1774 | 3325 |
| **rRNA** | | | | | | |
| unique reads | 3771 | 3643 | 3705 | 4236 | 2351 | 4295 |
| **median error rate** | | | | | | |
| complete reference | 7.5% | 8.5% | 7.7% | 8.0% | 8.9% | 7.4% |
| *T. b.* chromosomes | 7.5% | 8.6% | 7.8% | 8.0% | 9.0% | 7.5% |
| RCS | 4.9% | 5.9% | 5.2% | 5.2% | 5.9% | 4.6% |

| | | | | | | |
|---|---|---|---|---|---|---|
| maxicircle | 7.1% | 7.8% | 7.3% | 7.7% | 8.2% | 7.3% |
| **genome coverage** | | | | | | |
| depth > 0 | 64% | 67% | 67% | 73% | 65% | 67% |
| depth > 4 | 37% | 43% | 45% | 56% | 44% | 46% |
| **annotated exons (N=10728)** | | | | | | |
| depth > 0 | 8463 | 8551 | 8559 | 9018 | 8904 | 8899 |
| depth > 4 | 7049 | 7487 | 7664 | 8436 | 8187 | 8238 |

**Supplementary Table S3.** Identification of full-length DRS sequencing reads. Full-length reads were identified based on the presence of at least 15nt of the spliced leader (SL)-RNA sequence at the 5'-end (5'-SL) in addition to a 3'-poly(A) sequence identified by the poly(A) module of nanopolish. rep=DRS library replicates of procyclic-stage and bloodstream-stage *T. brucei*.

| | **procyclic-stage libraries** | | | **bloodstream-stage libraries** | | |
|---|---|---|---|---|---|---|
| | **rep1** | **rep2** | **rep3** | **rep1** | **rep2** | **rep3** |
| input | 2.59E+05 | 3.03E+05 | 3.35E+05 | 5.04E+05 | 5.24E+05 | 4.16E+05 |
| 5'-SL | 1.53E+05 | 1.73E+05 | 1.81E+05 | 1.86E+05 | 1.43E+05 | 1.32E+05 |
| | 59% | 57% | 54% | 37% | 27% | 32% |
| 3'-poly(A) | 2.24E+05 | 2.47E+05 | 2.85E+05 | 4.29E+05 | 4.33E+05 | 3.50E+05 |
| | 87% | 82% | 85% | 85% | 83% | 84% |
| 5'-SL & 3'-poly(A) | 1.37E+05 | 1.45E+05 | 1.57E+05 | 1.62E+05 | 1.22E+05 | 1.15E+05 |
| | 53% | 48% | 47% | 32% | 23% | 28% |

**Supplementary Table S4.** Performance of lncRNA prediction tools. DRS sequences (supported by minimally 3 reads) were analyzed for their coding potential using CPC2 (Kang et al., 2017) or LncFinder (Han et al., 2019). Genomic coordinates of sequences predicted as noncoding were compared to the genome annotation file and checked for overlap with the coding sequence (CDS) of annotated mRNAs.

| | **total** | **predicted as non-coding** | | |
|---|---|---|---|---|
| | | CPC2 | LncFinder | overlap |
| input | 10840 | 2513 | 1825 | 1820 |
| annotated CDS | 8496 | 434 | 99 | 96 |
| | | 17.3% | 5.4% | 5.3% |

| gene product | annotated | CPC2 | LncFinder | overlap |
|---|---|---|---|---|
| ribosomal protein | 254 | 38 | 1 | 0 |
| histone | 64 | 30 | 0 | 0 |
| dynein | 48 | 8 | 0 | 0 |
| hypothetical/unspecified | 2851 | 223 | 4 | 4 |

**Supplementary Table S5.** Genomic locations of novel intergenic lncRNAs. Overlapping transcripts are shaded. LncRNAs of similar sequence but localizing to a different genomic locus are given in the last column. ID=identification number. nt=novel transcript.

| transcript-ID | chromosome | start | end | strand | similar to or overlapping with |
|---|---|---|---|---|---|
| nt_194.4 | Tb927_01_v5.1 | 8677 | 9311 | - | |
| nt_200.16 | Tb927_01_v5.1 | 59156 | 59765 | - | |
| nt_200.17 | Tb927_01_v5.1 | 59366 | 59765 | - | |
| nt_202.1 | Tb927_01_v5.1 | 63008 | 64548 | - | |
| nt_205.1 | Tb927_01_v5.1 | 72570 | 74748 | - | |
| nt_1.2 | Tb927_01_v5.1 | 93131 | 94573 | + | |
| nt_212.5 | Tb927_01_v5.1 | 102280 | 103253 | - | |
| nt_213.1 | Tb927_01_v5.1 | 103403 | 103992 | - | |
| nt_215.1 | Tb927_01_v5.1 | 127544 | 128314 | - | |
| nt_10.2 | Tb927_01_v5.1 | 133736 | 134603 | + | nt_215.2 |
| nt_222.2 | Tb927_01_v5.1 | 174593 | 176674 | - | |
| nt_17.3 | Tb927_01_v5.1 | 188014 | 189511 | + | |
| nt_249.1 | Tb927_01_v5.1 | 279320 | 279778 | - | |
| nt_255.1 | Tb927_01_v5.1 | 643401 | 644549 | - | nt_256.1 |
| nt_256.1 | Tb927_01_v5.1 | 647417 | 648547 | - | nt_255.1 |
| nt_416.3 | Tb927_02_v5.1 | 4330 | 6766 | - | |
| nt_417.1 | Tb927_02_v5.1 | 6890 | 7248 | - | |
| nt_417.2 | Tb927_02_v5.1 | 6890 | 8350 | - | |
| nt_419.1 | Tb927_02_v5.1 | 27741 | 29774 | - | |
| nt_421.1 | Tb927_02_v5.1 | 32945 | 35103 | - | |
| nt_423.1 | Tb927_02_v5.1 | 38299 | 40726 | - | nt_430.1 |
| nt_430.1 | Tb927_02_v5.1 | 69265 | 71691 | - | nt_423.1 |
| nt_433.3 | Tb927_02_v5.1 | 79896 | 81446 | - | |
| nt_445.1 | Tb927_02_v5.1 | 114448 | 115421 | - | |
| nt_447.2 | Tb927_02_v5.1 | 135155 | 136402 | - | |
| nt_451.3 | Tb927_02_v5.1 | 147956 | 148930 | - | |
| nt_468.11 | Tb927_02_v5.1 | 226734 | 227355 | - | |
| nt_300.1 | Tb927_02_v5.1 | 250325 | 250814 | + | |
| nt_303.2 | Tb927_02_v5.1 | 257928 | 258927 | + | |
| nt_473.1 | Tb927_02_v5.1 | 274417 | 275641 | - | |
| nt_474.1 | Tb927_02_v5.1 | 301020 | 301646 | - | |
| nt_320.1 | Tb927_02_v5.1 | 358649 | 359954 | + | |
| nt_478.1 | Tb927_02_v5.1 | 384788 | 386406 | - | |
| nt_345.1 | Tb927_02_v5.1 | 600480 | 601455 | + | nt_350.1 |
| nt_345.2 | Tb927_02_v5.1 | 600881 | 601455 | + | nt_350.2, nt_350.1, nt_347.2 |
| nt_347.2 | Tb927_02_v5.1 | 604426 | 604975 | + | nt_345.2, nt_345.1, nt_350.1, nt_350.2 |
| nt_350.1 | Tb927_02_v5.1 | 611119 | 612056 | + | nt_345.1 |
| nt_350.2 | Tb927_02_v5.1 | 611510 | 612056 | + | nt_345.2, nt_345.1, nt_347.2 |
| nt_352.1 | Tb927_02_v5.1 | 614653 | 614930 | + | nt_350.1, nt_345.1 |
| nt_596.1 | Tb927_02_v5.1 | 979948 | 980333 | - | nt_375.1 |
| nt_599.1 | Tb927_02_v5.1 | 995887 | 996615 | - | nt_373.2 |
| nt_373.2 | Tb927_02_v5.1 | 998092 | 998818 | + | nt_599.1 |
| nt_373.3 | Tb927_02_v5.1 | 998635 | 1000193 | + | |
| nt_373.5 | Tb927_02_v5.1 | 998945 | 1000193 | + | |
| nt_375.1 | Tb927_02_v5.1 | 1014366 | 1014751 | + | nt_596.1 |
| nt_401.6 | Tb927_02_v5.1 | 1122445 | 1123351 | + | |
| nt_401.7 | Tb927_02_v5.1 | 1122445 | 1124737 | + | |
| nt_401.8 | Tb927_02_v5.1 | 1123050 | 1124737 | + | |
| nt_401.9 | Tb927_02_v5.1 | 1123464 | 1124737 | + | |
| nt_411.1 | Tb927_02_v5.1 | 1164676 | 1166191 | + | |
| nt_413.2 | Tb927_02_v5.1 | 1168338 | 1168836 | + | |
| nt_413.3 | Tb927_02_v5.1 | 1168338 | 1169283 | + | |
| nt_841.3 | Tb927_03_v5.1 | 384581 | 385789 | - | |
| nt_844.1 | Tb927_03_v5.1 | 391980 | 393761 | - | |
| nt_990.4 | Tb927_03_v5.1 | 1160150 | 1160561 | - | |
| nt_1160.1 | Tb927_04_v5.1 | 199 | 700 | - | |
| nt_1161.1 | Tb927_04_v5.1 | 815 | 1287 | - | |
| nt_1162.2 | Tb927_04_v5.1 | 7423 | 8867 | - | |
| nt_1162.3 | Tb927_04_v5.1 | 7888 | 8867 | - | |
| nt_1107.1 | Tb927_04_v5.1 | 692681 | 693120 | + | |

| | | | | | |
|---|---|---|---|---|---|
| nt_1399.1 | Tb927_04_v5.1 | 1172047 | 1172332 | - | |
| nt_1158.1 | Tb927_04_v5.1 | 1472777 | 1474585 | + | |
| nt_1158.4 | Tb927_04_v5.1 | 1474722 | 1476645 | + | |
| nt_1742.1 | Tb927_05_v5.1 | 13140 | 14681 | - | |
| nt_1742.3 | Tb927_05_v5.1 | 13807 | 14681 | - | |
| nt_1743.1 | Tb927_05_v5.1 | 15984 | 19513 | - | |
| nt_1744.1 | Tb927_05_v5.1 | 19648 | 20803 | - | |
| nt_1744.6 | Tb927_05_v5.1 | 20808 | 22315 | - | |
| nt_1487.1 | Tb927_05_v5.1 | 27791 | 28386 | + | |
| nt_1745.2 | Tb927_05_v5.1 | 66202 | 68362 | - | nt_1493.5, nt_1493.6 |
| nt_1750.1 | Tb927_05_v5.1 | 83729 | 84843 | - | nt_1762.2 |
| nt_1750.2 | Tb927_05_v5.1 | 83729 | 86415 | - | |
| nt_1489.2 | Tb927_05_v5.1 | 100971 | 101950 | + | |
| nt_1492.1 | Tb927_05_v5.1 | 108791 | 109331 | + | |
| nt_1493.1 | Tb927_05_v5.1 | 113480 | 114387 | + | |
| nt_1493.2 | Tb927_05_v5.1 | 113480 | 115188 | + | |
| nt_1493.3 | Tb927_05_v5.1 | 114523 | 115188 | + | |
| nt_1493.4 | Tb927_05_v5.1 | 114523 | 117496 | + | |
| nt_1493.5 | Tb927_05_v5.1 | 115337 | 117496 | + | |
| nt_1493.6 | Tb927_05_v5.1 | 116306 | 117496 | + | nt_1745.2 |
| nt_1759.1 | Tb927_05_v5.1 | 134729 | 135378 | - | |
| nt_1762.2 | Tb927_05_v5.1 | 146145 | 148907 | - | |
| nt_1763.1 | Tb927_05_v5.1 | 149009 | 149832 | - | |
| nt_1768.1 | Tb927_05_v5.1 | 169875 | 170775 | - | |
| nt_1768.2 | Tb927_05_v5.1 | 169875 | 171185 | - | |
| nt_1768.4 | Tb927_05_v5.1 | 169875 | 172240 | - | |
| nt_1768.7 | Tb927_05_v5.1 | 170931 | 172240 | - | |
| nt_1768.8 | Tb927_05_v5.1 | 171302 | 172240 | - | |
| nt_1500.1 | Tb927_05_v5.1 | 176139 | 176983 | + | |
| nt_1769.1 | Tb927_05_v5.1 | 184812 | 185584 | - | |
| nt_1504.1 | Tb927_05_v5.1 | 249055 | 249330 | + | |
| nt_1527.1 | Tb927_05_v5.1 | 353506 | 353925 | + | |
| nt_1667.1 | Tb927_05_v5.1 | 929096 | 929834 | + | |
| nt_1725.1 | Tb927_05_v5.1 | 1395730 | 1396781 | + | |
| nt_1728.1 | Tb927_05_v5.1 | 1409234 | 1410517 | + | |
| nt_1729.4 | Tb927_05_v5.1 | 1416865 | 1417172 | + | |
| nt_1737.1 | Tb927_05_v5.1 | 1435122 | 1436168 | + | |
| nt_1738.1 | Tb927_05_v5.1 | 1438285 | 1439317 | + | |
| nt_1922.3 | Tb927_06_v5.1 | 22976 | 23454 | + | |
| nt_1922.4 | Tb927_06_v5.1 | 22976 | 24015 | + | |
| nt_2127.1 | Tb927_06_v5.1 | 32116 | 32745 | - | |
| nt_2128.1 | Tb927_06_v5.1 | 32860 | 33654 | - | |
| nt_2129.1 | Tb927_06_v5.1 | 33779 | 34307 | - | nt_2132.4, nt_1935.1 |
| nt_1923.3 | Tb927_06_v5.1 | 38192 | 39606 | + | |
| nt_1923.5 | Tb927_06_v5.1 | 39808 | 41546 | + | nt_2143.1 |
| nt_2132.4 | Tb927_06_v5.1 | 77739 | 78283 | - | nt_1935.1, nt_2129.1 |
| nt_1928.1 | Tb927_06_v5.1 | 90325 | 91026 | + | nt_2136.1, nt_2136.3, nt_2140.2, nt_2140.1, nt_1940.3 |
| nt_2136.1 | Tb927_06_v5.1 | 117583 | 120484 | - | |
| nt_2136.3 | Tb927_06_v5.1 | 119616 | 120484 | - | nt_1928.1, nt_2140.1, nt_2140.2 |
| nt_2138.1 | Tb927_06_v5.1 | 125083 | 125343 | - | nt_2142.1 |
| nt_2140.1 | Tb927_06_v5.1 | 130027 | 131874 | - | nt_2136.1 |
| nt_2140.2 | Tb927_06_v5.1 | 130996 | 131874 | - | nt_2136.3, nt_2136.1, nt_1940.3, nt_1928.1 |
| nt_2140.3 | Tb927_06_v5.1 | 131458 | 131874 | - | |
| nt_2142.1 | Tb927_06_v5.1 | 136469 | 136723 | - | nt_2138.1 |
| nt_2143.1 | Tb927_06_v5.1 | 141355 | 143137 | - | nt_1923.5 |
| nt_1935.1 | Tb927_06_v5.1 | 148639 | 149183 | + | nt_2132.4 |
| nt_1937.1 | Tb927_06_v5.1 | 150201 | 150812 | + | |
| nt_1940.2 | Tb927_06_v5.1 | 160294 | 161157 | + | |
| nt_1965.1 | Tb927_06_v5.1 | 339689 | 340081 | + | |
| nt_2178.1 | Tb927_06_v5.1 | 363244 | 363619 | - | |
| nt_1983.1 | Tb927_06_v5.1 | 728981 | 729771 | + | |
| nt_2374.1 | Tb927_07_v5.1 | 32685 | 34046 | + | |
| nt_2707.1 | Tb927_07_v5.1 | 479102 | 479709 | - | nt_2710.1, nt_2713.1 |
| nt_2710.1 | Tb927_07_v5.1 | 484661 | 485296 | - | nt_2707.1, nt_2713.1 |
| nt_2713.1 | Tb927_07_v5.1 | 501461 | 502096 | - | nt_2710.1, nt_2707.1 |
| nt_2726.4 | Tb927_07_v5.1 | 543327 | 543837 | - | |
| nt_2496.1 | Tb927_07_v5.1 | 834988 | 836167 | + | |
| nt_2856.1 | Tb927_07_v5.1 | 1238481 | 1239149 | - | |

| | | | | | |
|---|---|---|---|---|---|
| nt_2875.1 | Tb927_07_v5.1 | 1285552 | 1286087 | - | |
| nt_2881.1 | Tb927_07_v5.1 | 1308199 | 1309532 | - | |
| nt_2615.1 | Tb927_07_v5.1 | 1601963 | 1602373 | + | |
| nt_2633.1 | Tb927_07_v5.1 | 1790335 | 1790900 | + | |
| nt_2926.1 | Tb927_07_v5.1 | 1839125 | 1839473 | - | |
| nt_2983.1 | Tb927_07_v5.1 | 2096938 | 2097338 | - | |
| nt_2983.3 | Tb927_07_v5.1 | 2096947 | 2097338 | - | |
| nt_3001.1 | Tb927_08_v5.1 | 49746 | 50574 | + | |
| nt_3446.1 | Tb927_08_v5.1 | 296504 | 297024 | - | |
| nt_3073.1 | Tb927_08_v5.1 | 791866 | 792596 | + | |
| nt_3089.1 | Tb927_08_v5.1 | 856518 | 857230 | + | |
| nt_3144.4 | Tb927_08_v5.1 | 1045706 | 1046768 | + | |
| nt_3511.1 | Tb927_08_v5.1 | 1056525 | 1057061 | - | |
| nt_3541.1 | Tb927_08_v5.1 | 1155854 | 1157057 | - | |
| nt_3260.1 | Tb927_08_v5.1 | 1998335 | 1999109 | + | |
| nt_3316.1 | Tb927_08_v5.1 | 2201558 | 2201957 | + | nt_3320.1, nt_3318.2 |
| nt_3318.2 | Tb927_08_v5.1 | 2204283 | 2204682 | + | nt_3320.1, nt_3316.1 |
| nt_3320.1 | Tb927_08_v5.1 | 2207008 | 2207407 | + | nt_3318.2, nt_3316.1 |
| nt_3670.1 | Tb927_08_v5.1 | 2236556 | 2237618 | - | |
| nt_3396.1 | Tb927_08_v5.1 | 2459446 | 2459948 | + | |
| nt_4004.1 | Tb927_09_v5.1 | 158853 | 159499 | - | |
| nt_4020.1 | Tb927_09_v5.1 | 351703 | 352172 | - | |
| nt_4082.1 | Tb927_09_v5.1 | 613931 | 614541 | - | nt_4099.1, nt_4091.1, nt_4089.1 |
| nt_4089.1 | Tb927_09_v5.1 | 626134 | 626762 | - | nt_4091.1, nt_4099.1, nt_4082.1 |
| nt_4091.1 | Tb927_09_v5.1 | 629231 | 629858 | - | nt_4089.1, nt_4099.1, nt_4082.1 |
| nt_4099.1 | Tb927_09_v5.1 | 648586 | 649212 | - | nt_4082.1, nt_4089.1, nt_4091.1 |
| nt_4118.1 | Tb927_09_v5.1 | 1189203 | 1189814 | - | nt_4124.3 |
| nt_4120.1 | Tb927_09_v5.1 | 1192799 | 1195231 | - | |
| nt_4120.2 | Tb927_09_v5.1 | 1194028 | 1195231 | - | |
| nt_4124.3 | Tb927_09_v5.1 | 1211124 | 1211735 | - | nt_4118.1 |
| nt_4146.1 | Tb927_09_v5.1 | 1303364 | 1304868 | - | |
| nt_4220.1 | Tb927_09_v5.1 | 1525948 | 1526948 | - | |
| nt_4243.7 | Tb927_09_v5.1 | 1611151 | 1611571 | - | |
| nt_3837.1 | Tb927_09_v5.1 | 1824285 | 1824705 | + | |
| nt_4296.1 | Tb927_09_v5.1 | 2087164 | 2088287 | - | |
| nt_3936.1 | Tb927_09_v5.1 | 2286546 | 2287343 | + | |
| nt_3972.4 | Tb927_09_v5.1 | 2440372 | 2440616 | + | nt_3975.1 |
| nt_3975.1 | Tb927_09_v5.1 | 2443945 | 2444189 | + | nt_3972.4 |
| nt_3976.5 | Tb927_09_v5.1 | 2445730 | 2446145 | + | |
| nt_3977.3 | Tb927_09_v5.1 | 2448087 | 2448768 | + | |
| nt_4363.1 | Tb927_09_v5.1 | 2512687 | 2513703 | - | |
| nt_4363.2 | Tb927_09_v5.1 | 2512687 | 2515347 | - | |
| nt_3987.6 | Tb927_09_v5.1 | 2523052 | 2524193 | + | |
| nt_4377.1 | Tb927_09_v5.1 | 2668064 | 2669227 | - | |
| nt_4381.1 | Tb927_09_v5.1 | 2678157 | 2678924 | - | |
| nt_3994.1 | Tb927_09_v5.1 | 2690171 | 2690831 | + | |
| nt_3995.1 | Tb927_09_v5.1 | 2692919 | 2694243 | + | |
| nt_3996.1 | Tb927_09_v5.1 | 2696153 | 2697089 | + | |
| nt_4397.1 | Tb927_09_v5.1 | 3405122 | 3406513 | - | |
| nt_5045.1 | Tb927_10_v5.1 | 21824 | 23765 | - | |
| nt_5049.1 | Tb927_10_v5.1 | 37614 | 38106 | - | |
| nt_4426.1 | Tb927_10_v5.1 | 140102 | 140474 | + | |
| nt_4456.1 | Tb927_10_v5.1 | 257007 | 257389 | + | |
| nt_5077.1 | Tb927_10_v5.1 | 365551 | 366118 | - | |
| nt_5083.1 | Tb927_10_v5.1 | 391955 | 392513 | - | |
| nt_5089.2 | Tb927_10_v5.1 | 425194 | 425582 | - | |
| nt_4462.1 | Tb927_10_v5.1 | 452242 | 453144 | + | |
| nt_5101.1 | Tb927_10_v5.1 | 470525 | 471426 | - | |
| nt_4504.1 | Tb927_10_v5.1 | 580560 | 581061 | + | |
| nt_4515.1 | Tb927_10_v5.1 | 612769 | 613390 | + | |
| nt_4554.1 | Tb927_10_v5.1 | 745710 | 746110 | + | |
| nt_4561.6 | Tb927_10_v5.1 | 777032 | 778057 | + | |
| nt_4580.1 | Tb927_10_v5.1 | 837019 | 837242 | + | |
| nt_4638.3 | Tb927_10_v5.1 | 1289793 | 1290560 | + | |
| nt_4641.4 | Tb927_10_v5.1 | 1302543 | 1303880 | + | |
| nt_4642.1 | Tb927_10_v5.1 | 1304003 | 1304550 | + | |
| nt_5222.1 | Tb927_10_v5.1 | 1493180 | 1493729 | - | |
| nt_5229.4 | Tb927_10_v5.1 | 1527472 | 1529007 | - | |
| nt_4701.3 | Tb927_10_v5.1 | 1684401 | 1685118 | + | |

| | | | | | |
|---|---|---|---|---|---|
| nt_4721.4 | Tb927_10_v5.1 | 1731709 | 1732213 | + | |
| nt_4739.1 | Tb927_10_v5.1 | 1787996 | 1788361 | + | |
| nt_5285.4 | Tb927_10_v5.1 | 1885770 | 1886833 | - | |
| nt_5285.5 | Tb927_10_v5.1 | 1885770 | 1887510 | - | |
| nt_5302.1 | Tb927_10_v5.1 | 1944645 | 1945114 | - | |
| nt_5319.8 | Tb927_10_v5.1 | 1999712 | 2000212 | - | |
| nt_4823.1 | Tb927_10_v5.1 | 2334878 | 2336931 | + | |
| nt_4829.2 | Tb927_10_v5.1 | 2349525 | 2350028 | + | |
| nt_5373.4 | Tb927_10_v5.1 | 2639414 | 2641305 | - | |
| nt_5373.5 | Tb927_10_v5.1 | 2639674 | 2641305 | - | |
| nt_5373.7 | Tb927_10_v5.1 | 2640254 | 2641305 | - | |
| nt_5374.1 | Tb927_10_v5.1 | 2641396 | 2642378 | - | |
| nt_5374.2 | Tb927_10_v5.1 | 2641396 | 2642574 | - | |
| nt_5387.1 | Tb927_10_v5.1 | 2688192 | 2688712 | - | |
| nt_4896.11 | Tb927_10_v5.1 | 2828919 | 2829983 | + | |
| nt_5425.1 | Tb927_10_v5.1 | 3027811 | 3028219 | - | |
| nt_5425.2 | Tb927_10_v5.1 | 3027811 | 3028626 | - | |
| nt_5425.3 | Tb927_10_v5.1 | 3027866 | 3028626 | - | |
| nt_5504.1 | Tb927_10_v5.1 | 3302765 | 3303306 | - | |
| nt_5600.9 | Tb927_10_v5.1 | 3616215 | 3616803 | - | |
| nt_5601.1 | Tb927_10_v5.1 | 3616907 | 3617888 | - | |
| nt_5601.2 | Tb927_10_v5.1 | 3616907 | 3619154 | - | |
| nt_5601.4 | Tb927_10_v5.1 | 3618107 | 3619154 | - | |
| nt_4981.3 | Tb927_10_v5.1 | 3724065 | 3724420 | + | |
| nt_5003.1 | Tb927_10_v5.1 | 3813154 | 3813738 | + | |
| nt_5013.1 | Tb927_10_v5.1 | 3865755 | 3866225 | + | |
| nt_5027.3 | Tb927_10_v5.1 | 3924719 | 3925126 | + | |
| nt_5625.2 | Tb927_11_RH_fork_v5.1 | 136516 | 137375 | - | |
| nt_5630.1 | Tb927_11_RH_fork_v5.1 | 237731 | 238549 | - | |
| nt_5631.1 | Tb927_11_RH_fork_v5.1 | 617151 | 618126 | - | |
| nt_6411.1 | Tb927_11_v5.1 | 29046 | 30954 | - | nt_5697.3 |
| nt_5691.2 | Tb927_11_v5.1 | 421251 | 422132 | + | |
| nt_5697.1 | Tb927_11_v5.1 | 444818 | 445778 | + | |
| nt_5697.2 | Tb927_11_v5.1 | 444818 | 446739 | + | |
| nt_6484.6 | Tb927_11_v5.1 | 551680 | 552075 | - | |
| nt_6484.7 | Tb927_11_v5.1 | 551680 | 552602 | - | |
| nt_6484.8 | Tb927_11_v5.1 | 552195 | 552602 | - | |
| nt_6556.1 | Tb927_11_v5.1 | 917913 | 918913 | - | |
| nt_5872.4 | Tb927_11_v5.1 | 1404891 | 1405479 | + | |
| nt_6609.4 | Tb927_11_v5.1 | 1751495 | 1751803 | - | |
| nt_6689.1 | Tb927_11_v5.1 | 2023993 | 2025461 | - | |
| nt_5965.4 | Tb927_11_v5.1 | 2046931 | 2047701 | + | |
| nt_5989.1 | Tb927_11_v5.1 | 2110152 | 2110889 | + | |
| nt_6236.1 | Tb927_11_v5.1 | 3149663 | 3150319 | + | |
| nt_6773.2 | Tb927_11_v5.1 | 3206614 | 3208824 | - | |
| nt_6773.12 | Tb927_11_v5.1 | 3209915 | 3210568 | - | |
| nt_6256.1 | Tb927_11_v5.1 | 3214193 | 3215719 | + | |
| nt_6258.1 | Tb927_11_v5.1 | 3216967 | 3217711 | + | |
| nt_6258.2 | Tb927_11_v5.1 | 3216967 | 3218598 | + | |
| nt_6807.1 | Tb927_11_v5.1 | 3515911 | 3517185 | - | |
| nt_6809.1 | Tb927_11_v5.1 | 3521361 | 3522957 | - | |

**Supplementary Table S6.** Differential gene expression. Genes up-regulated in bloodstream-stage trypanosomes.

| gene-ID | fold change (BS/PC) | adjusted *p*-value | gene product |
|---|---|---|---|
| Tb927.9.16490 | 143.20 | 7.21E-23 | variant surface glycoprotein (VSG), putative |
| Tb927.1.20 | 132.70 | 2.47E-21 | expression site-associated gene 3 (ESAG3), pseudogene |
| Tb927.1.5100 | 97.18 | 2.82E-16 | expression site-associated gene 2 (ESAG2) protein, putative |
| Tb927.1.4910 | 95.78 | 1.59E-16 | expression site-associated gene 1 (ESAG1) protein, putative |
| Tb927.5.150 | 79.35 | 3.12E-13 | hypothetical protein, conserved |
| Tb927.2.3320 | 75.78 | 8.34E-13 | 65 kDa invariant surface glycoprotein |
| Tb11.01.6240 | 71.05 | 9.28E-12 | expression site-associated gene 2 (ESAG2) protein, putative |
| nt_5631.1 | 70.64 | 1.43E-11 | lncRNA, putative |
| Tb11.01.6250 | 69.34 | 3.92E-11 | expression site-associated gene 11 (ESAG11) protein, putative |
| Tb927.7.170 | 69.15 | 6.91E-11 | expression site-associated gene 9 (ESAG9) protein, putative |
| Tb927.1.5110 | 68.85 | 1.67E-11 | expression site-associated gene 11 (ESAG11) protein, putative |
| Tb927.9.7380 | 68.60 | 2.54E-11 | variant surface glycoprotein (VSG)-related, putative |
| Tb927.5.1390 | 66.47 | 8.13E-11 | 64 kDa invariant surface glycoprotein |
| Tb927.5.120 | 66.34 | 1.57E-10 | expression site-associated gene 9 (ESAG9) protein, putative |
| Tb927.2.6180 | 66.20 | 2.50E-10 | iron/ascorbate oxidoreductase family protein, putative |
| Tb927.1.4870 | 65.73 | 1.33E-10 | expression site-associated gene 1 (ESAG1) protein, putative |
| Tb927.9.7320 | 64.18 | 6.15E-10 | expression site-associated gene 11 (ESAG11) protein, putative |
| Tb927.1.4900 | 64.07 | 1.28E-18 | expression site-associated gene 11 (ESAG11) protein, putative |
| Tb927.9.16880 | 63.67 | 4.01E-10 | expression site-associated gene 3 (ESAG3, pseudogene), putative |
| Tb927.9.16500 | 63.22 | 3.16E-09 | variant surface glycoprotein (VSG, atypical), putative |
| Tb927.11.4100 | 63.06 | 5.74E-10 | variant surface glycoprotein (VSG), putative |
| nt_194.4 | 62.33 | 1.04E-09 | lncRNA, putative |
| Tb927.3.570 | 61.02 | 2.87E-09 | expression site-associated gene 2 (ESAG2) protein, putative |
| KS17gene_223a | 61.01 | 1.37E-09 | lncRNA, putative |
| Tb927.3.1520 | 60.61 | 1.34E-08 | variant surface glycoprotein (VSG)-related, putative |
| Tb927.2.3310 | 59.88 | 2.43E-09 | 65 kDa invariant surface glycoprotein |
| Tb927.3.560 | 59.36 | 5.15E-08 | expression site-associated gene 11 (ESAG11) protein, putative |
| Tb927.5.1400 | 58.96 | 1.33E-08 | hypothetical protein |
| Tb927.5.4900 | 58.55 | 4.22E-07 | variant surface glycoprotein, frameshift |
| Tb927.7.160 | 58.27 | 9.32E-09 | expression site-associated gene 2 (ESAG2), degenerate |
| Tb927.11.14610 | 57.96 | 1.49E-08 | procyclin-associated gene 4 (PAG4) protein, putative |
| Tb927.1.5120 | 57.53 | 1.11E-08 | expression site-associated gene 1 (ESAG1) protein, putative |
| Tb927.1.5240 | 56.62 | 1.96E-08 | expression site-associated gene 1 (ESAG1) protein, putative |
| nt_5045.1 | 56.58 | 1.64E-08 | lncRNA, putative |
| Tb927.2.6320 | 56.50 | 2.66E-08 | adenosine transporter 2, putative |
| Tb927.2.3295 | 55.98 | 5.53E-08 | unspecified product |
| Tb927.8.7330 | 55.59 | 3.91E-08 | hypothetical protein |
| Tb927.11.14620 | 55.17 | 9.38E-17 | expression site-associated gene 2 (ESAG2) protein, putative |
| KS17gene_225a | 54.70 | 4.96E-08 | lncRNA, putative |
| Tb927.3.1470 | 53.95 | 7.93E-08 | variant surface glycoprotein (VSG)-related, putative |
| nt_4829.2 | 53.75 | 9.41E-08 | lncRNA, putative |
| Tb927.9.7340 | 52.73 | 2.04E-07 | expression site-associated gene 9 (ESAG9) protein, putative |
| Tb927.5.1410 | 52.09 | 9.16E-07 | 64 kDa invariant surface glycoprotein |
| Tb927.3.5790 | 50.15 | 6.84E-07 | expression site-associated gene 9 (ESAG9) (pseudogene) |
| KS17gene_6206a | 50.10 | 1.90E-06 | lncRNA, putative |
| Tb927.3.5830 | 50.07 | 7.94E-07 | expression site-associated gene 1 (ESAG1) protein, putative |
| Tb927.9.7410 | 49.98 | 1.34E-06 | expression site-associated gene 2 (ESAG2) protein, putative |
| Tb927.11.14600 | 49.96 | 1.27E-06 | procyclin-associated gene 2-like protein, putative |
| Tb927.5.5450 | 49.92 | 7.96E-07 | Variant Surface Glycoprotein, putative |
| nt_1742.1 | 49.75 | 8.78E-07 | lncRNA, putative |
| KS17gene_2863a | 49.46 | 1.00E-06 | lncRNA, putative |
| Tb927.3.1500 | 49.14 | 1.79E-06 | variant surface glycoprotein (VSG)-related, putative |
| Tb927.11.18660 | 48.96 | 1.19E-06 | expression site-associated gene 9 (ESAG9), putative (pseudogene) |
| Tb927.9.730 | 48.87 | 1.54E-06 | hypothetical protein, conserved |
| nt_347.2 | 48.65 | 2.07E-06 | lncRNA, putative |
| Tb927.2.3315 | 48.63 | 6.93E-06 | unspecified product |
| Tb927.10.1780 | 48.23 | 3.59E-06 | hypothetical protein |
| Tb927.3.5820 | 48.17 | 2.41E-06 | expression site-associated gene 11 (ESAG11), degenerate |

| | | | |
|---|---|---|---|
| Tb927.10.5680 | 47.98 | 2.00E-06 | procyclin-associated gene 1 (PAG1) protein, putative |
| Tb927.3.2590 | 47.92 | 2.22E-06 | hypothetical protein |
| Tb927.5.1420 | 46.99 | 3.98E-06 | hypothetical protein |
| KS17gene_3556a | 46.91 | 4.01E-06 | lncRNA, putative |
| KS17gene_3552a | 46.64 | 3.81E-06 | lncRNA, putative |
| Tb09_snoRNA_0076 | 46.30 | 8.06E-06 | H/ACA snoRNA, TB9Cs2H1 |
| Tb927.9.2855 | 45.64 | 7.25E-06 | Domain of unknown function (DUF5075), putative |
| nt_844.1 | 45.51 | 9.08E-06 | lncRNA, putative |
| Tb927.2.6230 | 45.25 | 7.82E-06 | iron/ascorbate oxidoreductase family protein, putative |
| Tb927.8.1665 | 45.16 | 9.22E-06 | hypothetical protein |
| Tb927.1.4890 | 45.07 | 1.14E-13 | expression site-associated gene 2 (ESAG2) protein, putative |
| Tb927.6.1340 | 44.80 | 1.25E-05 | cyclophilin-type peptidyl-prolyl cis-trans isomerase, putative |
| Tb927.2.200 | 44.28 | 4.05E-18 | expression site-associated gene 3 (ESAG3), degenerate |
| Tb927.3.5810 | 44.22 | 1.10E-05 | expression site-associated gene 2 (ESAG2) (pseudogene) |
| Tb927.3.520 | 44.15 | 1.59E-05 | expression site-associated gene 1 (ESAG1) protein, putative |
| KS17gene_943a | 42.59 | 2.39E-05 | lncRNA, putative |
| Tb927.1.5170 | 41.74 | 4.45E-05 | variant surface glycoprotein (VSG)-related, putative |
| Tb927.11.12120 | 41.53 | 4.71E-05 | RNA-binding protein, putative |
| Tb927.5.110 | 41.47 | 3.68E-05 | variant surface glycoprotein (VSG)-related, putative |
| KS17gene_6881a | 41.26 | 7.86E-05 | lncRNA, putative |
| Tb927.6.2520 | 41.03 | 5.23E-05 | hypothetical protein, conserved |
| Tb927.7.3260 | 40.63 | 9.12E-17 | expression site-associated gene 7 (ESAG7) protein, putative |
| Tb927.5.130 | 40.20 | 1.15E-04 | variant surface glycoprotein (VSG)-related, putative |
| Tb927.3.5800 | 39.77 | 1.29E-04 | expression site-associated gene 1 (ESAG1), degenerate |
| Tb927.2.3270 | 39.38 | 1.11E-17 | 65 kDa invariant surface glycoprotein |
| Tb927.5.1430 | 38.82 | 1.30E-04 | 64 kDa invariant surface glycoprotein |
| nt_4004.1 | 38.77 | 1.06E-04 | lncRNA, putative |
| nt_350.1 | 37.98 | 2.55E-04 | lncRNA, putative |
| Tb927.2.3280 | 37.74 | 3.01E-04 | 65 kDa invariant surface glycoprotein |
| nt_401.7 | 37.29 | 2.07E-04 | lncRNA, putative |
| Tb927.11.18650 | 36.22 | 4.15E-04 | expression site-associated gene 1 (ESAG1) protein, putative |
| Tb927.2.6220 | 35.81 | 4.12E-04 | adenosine transporter 2, putative |
| Tb927.9.7350 | 35.36 | 4.30E-04 | expression site-associated gene 1 (ESAG1), pseudogene |
| Tb927.6.1390 | 35.21 | 4.61E-04 | Trypanosomal VSG domain containing protein, putative |
| KS17gene_4587a | 34.85 | 5.86E-04 | lncRNA, putative |
| Tb927.9.7290 | 33.63 | 7.73E-04 | variant surface glycoprotein (VSG)-related, putative |
| nt_2633.1 | 33.55 | 8.45E-04 | lncRNA, putative |
| Tb927.2.3290 | 33.51 | 2.27E-15 | 65 kDa invariant surface glycoprotein |
| KS17gene_202a | 32.37 | 1.16E-03 | lncRNA, putative |
| KS17gene_4541a | 32.37 | 1.11E-03 | lncRNA, putative |
| Tb927.7.7510 | 32.35 | 1.63E-24 | hypothetical protein |
| Tb927.3.550 | 31.71 | 1.47E-03 | expression site-associated gene 1 (ESAG1), degenerate |
| KS17gene_7015a | 31.59 | 9.15E-13 | lncRNA, putative |
| Tb927.2.1380 | 31.33 | 1.57E-09 | leucine-rich repeat protein (LRRP), putative |
| Tb927.6.340 | 31.15 | 1.71E-03 | receptor-type adenylate cyclase GRESAG 4, pseudogene, putative |
| Tb927.2.2029 | 30.96 | 1.93E-09 | expression site-associated gene 3 (ESAG3), degenerate |
| nt_345.1 | 30.42 | 2.11E-03 | lncRNA, putative |
| Tb927.7.3250 | 30.34 | 9.81E-15 | expression site-associated gene 6 (ESAG6) protein, putative |
| Tb927.4.230 | 30.21 | 4.18E-09 | DNA-directed RNA polymerase III subunit, pseudogene, putative |
| Tb927.5.309b | 30.14 | 2.22E-12 | invariant surface glycoprotein, putative |
| KS17gene_215a | 28.57 | 9.06E-12 | lncRNA, putative |
| Tb927.10.1480 | 28.39 | 2.61E-08 | hypothetical protein |
| Tb927.10.9450 | 27.25 | 4.98E-13 | invariant surface glycoprotein, putative |
| Tb927.11.17890 | 26.97 | 6.95E-13 | expression site-associated gene 1 (ESAG1) protein, putative |
| Tb927.10.5710 | 26.14 | 1.88E-12 | hypothetical protein, conserved |
| Tb927.5.400 | 26.09 | 1.64E-10 | 75 kDa invariant surface glycoprotein, putative |
| Tb927.4.200 | 26.00 | 1.24E-07 | retrotransposon hot spot protein 1 (RHS1), putative |
| Tb927.6.1040 | 25.35 | 7.92E-15 | cysteine peptidase, Clan CA, family C1, Cathepsin L-like |
| Tb927.6.5180 | 25.09 | 3.68E-15 | retrotransposon hot spot protein 4 (RHS4), interrupted |
| Tb927.1.4860 | 24.90 | 4.49E-07 | expression site-associated gene 11 (ESAG11), pseudogene |
| Tb927.4.1230 | 24.38 | 4.77E-17 | hypothetical protein |
| nt_4823.1 | 24.32 | 5.15E-07 | lncRNA, putative |
| Tb927.10.12800 | 23.88 | 1.09E-08 | Zinc finger CCCH domain-containing protein 38 |

| | | | |
|---|---|---|---|
| Tb927.9.16050 | 23.29 | 1.39E-06 | leucine-rich repeat protein (pseudogene), putative |
| Tb927.7.180 | 22.61 | 1.14E-05 | Trypanosomal VSG domain containing protein, putative |
| KS17gene_5352a | 21.57 | 6.56E-11 | lncRNA, putative |
| Tb927.5.160 | 21.44 | 5.25E-06 | retrotransposon hot spot protein (RHS, pseudogene), putative |
| KS17gene_7014a | 21.22 | 1.08E-05 | lncRNA, putative |
| KS17gene_265a | 21.12 | 6.56E-11 | lncRNA, putative |
| Tb927.8.6760 | 20.76 | 9.23E-15 | translationally-controlled tumor protein homolog, putative |
| Tb927.10.5690 | 20.67 | 2.38E-08 | procyclin-associated gene 2 (PAG2) protein, putative |
| Tb927.6.1000 | 20.05 | 6.01E-16 | cysteine peptidase, Clan CA, family C1, Cathepsin L-like |
| Tb927.11.2400 | 19.73 | 1.90E-07 | Flabarin-like protein |
| Tb927.2.3340 | 19.45 | 9.43E-08 | hypothetical protein |
| nt_3987.6 | 18.58 | 1.20E-07 | lncRNA, putative |
| Tb927.11.4120 | 18.50 | 5.35E-05 | leucine-rich repeat protein (LRRP), putative |
| Tb927.2.980 | 18.40 | 3.51E-07 | retrotransposon hot spot protein 5 (RHS5), degenerate |
| Tb927.10.9510 | 18.19 | 8.66E-05 | hypothetical protein |
| KS17gene_212a | 18.10 | 7.64E-05 | lncRNA, putative |
| Tb927.1.5060 | 17.86 | 1.53E-08 | variant surface glycoprotein (VSG)-related, putative |
| KS17gene_4915a | 17.77 | 1.27E-04 | lncRNA, putative |
| Tb927.1.5200 | 17.68 | 9.83E-07 | expression site-associated gene 1 (ESAG1) protein, putative |
| Tb927.9.15680 | 17.49 | 1.03E-04 | expression site-associated gene 6 (ESAG6) |
| Tb927.10.10360 | 17.15 | 8.24E-07 | Microtubule-associated repetitive protein |
| nt_200.17 | 16.85 | 9.44E-07 | lncRNA, putative |
| Tb927.6.140 | 16.68 | 4.00E-12 | retrotransposon hot spot protein 5 (RHS5), putative |
| Tb927.3.4070 | 16.67 | 5.16E-18 | Pyruvate transporter, putative |
| nt_1487.1 | 16.39 | 3.40E-08 | lncRNA, putative |
| Tb927.1.2040 | 15.84 | 3.48E-04 | expression site-associated gene 2 (ESAG2) protein, putative |
| Tb927.6.330 | 15.78 | 8.88E-06 | receptor-type adenylate cyclase GRESAG 4, putative |
| Tb927.11.17880 | 15.66 | 4.19E-04 | expression site-associated gene 9 (ESAG9), degenerate |
| Tb927.9.15940 | 15.66 | 5.02E-04 | expression site-associated gene 3 (ESAG3) protein, putative |
| Tb927.7.370 | 15.62 | 4.41E-13 | hypothetical protein, conserved |
| KS17gene_2525a | 15.53 | 4.96E-04 | lncRNA, putative |
| Tb927.4.5260 | 15.27 | 1.65E-05 | UDP-Gal or UDP-GlcNAc-dependent glycosyltransferase, putative |
| Tb927.10.9465 | 15.23 | 1.36E-05 | hypothetical protein |
| Tb927.3.5690 | 15.13 | 8.84E-08 | hypothetical protein, conserved |
| Tb927.6.990 | 14.97 | 1.49E-14 | cysteine peptidase, Clan CA, family C1, Cathepsin L-like |
| Tb927.1.700 | 14.89 | 1.94E-16 | phosphoglycerate kinase |
| Tb927.4.4860 | 14.75 | 9.63E-16 | amino acid transporter 8, putative |
| KS17gene_4604a | 14.17 | 5.84E-12 | lncRNA, putative |
| Tb927.6.1050 | 14.07 | 4.51E-14 | cysteine peptidase, Clan CA, family C1, Cathepsin L-like |
| Tb927.5.293b | 14.05 | 5.61E-12 | hypothetical protein |
| Tb927.2.960 | 13.93 | 9.09E-09 | hypothetical protein |
| Tb927.7.6060 | 13.89 | 1.16E-03 | receptor-type adenylate cyclase GRESAG 4, putative |
| nt_4120.1 | 13.87 | 1.45E-03 | lncRNA, putative |
| Tb927.6.210 | 13.85 | 1.53E-03 | leucine-rich repeat protein (LRRP, pseudogene), putative |
| KS17gene_4363a | 13.65 | 1.36E-03 | lncRNA, putative |
| KS17gene_6546a | 13.57 | 1.52E-03 | lncRNA, putative |
| nt_447.2 | 13.04 | 2.34E-03 | lncRNA, putative |
| KS17gene_4610a | 12.97 | 1.40E-04 | lncRNA, putative |
| Tb927.11.11975 | 12.93 | 5.24E-15 | cytoskeleton-associated protein, putative |
| Tb927.1.430 | 12.92 | 1.28E-04 | retrotransposon hot spot protein (RHS, pseudogene), putative |
| Tb927.3.600 | 12.85 | 5.15E-11 | hypothetical protein |
| Tb927.5.4600 | 12.83 | 8.06E-09 | expression site-associated gene 3 (ESAG3) protein, putative |
| Tb927.2.490 | 12.77 | 4.40E-10 | DNA-directed RNA polymerase, pseudogene, putative |
| Tb8.NT.18 | 12.39 | 3.46E-03 | lncRNA, putative |
| Tb927.9.10650 | 12.02 | 9.09E-09 | hypothetical protein |
| Tb927.7.5790 | 11.92 | 2.16E-10 | protein disulfide isomerase, putative |
| Tb927.11.13630 | 11.90 | 1.17E-05 | hypothetical protein, conserved |
| Tb927.11.8490 | 11.77 | 2.15E-04 | DNA polymerase kappa, putative |
| Tb927.2.1330 | 11.59 | 2.16E-14 | retrotransposon hot spot protein 6 (RHS6), degenerate |
| Tb927.5.340 | 11.54 | 2.46E-06 | expression site-associated gene 5 (ESAG5) protein, putative |
| Tb927.4.4780 | 11.53 | 2.82E-09 | hypothetical protein |
| Tb927.8.3720 | 11.37 | 8.72E-10 | SUMO-interacting motif-containing protein |
| Tb927.9.7270 | 11.35 | 1.57E-06 | hypothetical protein |

| | | | |
|---|---|---|---|
| KS17gene_7564a | 11.28 | 5.81E-03 | lncRNA, putative |
| Tb927.1.1740 | 11.23 | 2.99E-04 | Microtubule-associated protein futsch, putative |
| nt_1492.1 | 11.12 | 3.67E-04 | lncRNA, putative |
| Tb927.7.7520 | 10.95 | 1.36E-15 | receptor-type adenylate cyclase GRESAG 4, putative |
| KS17gene_6494a | 10.91 | 6.62E-14 | lncRNA, putative |
| Tb927.6.350 | 10.68 | 5.89E-04 | hypothetical protein, conserved |
| Tb927.9.660 | 10.59 | 8.70E-03 | expression site-associated gene 1 (ESAG1) protein |
| Tb927.11.12130 | 10.55 | 8.51E-03 | hypothetical protein |
| Tb927.2.1110 | 10.48 | 4.80E-08 | DNA-directed RNA polymerase III subunit 2, pseudogene, putative |
| Tb927.8.5465 | 10.47 | 1.17E-10 | flagellar calcium-binding 24 kDa protein |
| Tb927.2.2020 | 10.35 | 3.23E-06 | expression site-associated gene 3 (ESAG3) protein, putative |
| Tb927.4.4870 | 10.32 | 2.54E-14 | amino acid transporter, putative |
| Tb927.9.740 | 10.23 | 5.55E-06 | hypothetical protein, conserved |
| Tb927.4.4450 | 10.13 | 9.29E-09 | adenylyl cyclase |
| Tb927.7.6500 | 10.12 | 1.68E-04 | variant surface glycoprotein (VSG), putative |
| Tb927.8.4110 | 10.01 | 8.47E-11 | Flagellum adhesion protein 3, putative |
| Tb927.8.1945 | 9.89 | 2.09E-10 | hypothetical protein |
| nt_10.2 | 9.83 | 1.24E-02 | lncRNA, putative |
| nt_1743.1 | 9.82 | 2.38E-07 | lncRNA, putative |
| Tb927.2.420 | 9.82 | 2.03E-13 | DNA-directed RNA polymerase, pseudogene, putative |
| Tb927.11.7550 | 9.79 | 2.81E-11 | hypothetical protein, conserved |
| Tb927.10.14890 | 9.73 | 2.48E-14 | C-terminal motor kinesin, putative |
| Tb927.5.4630 | 9.63 | 1.68E-04 | expression site-associated gene 1 (ESAG1) protein, putative |
| Tb927.5.4570 | 9.62 | 1.40E-06 | Flagellum adhesion protein 3 |
| Tb927.2.540 | 9.62 | 1.06E-13 | DNA-directed RNA polymerase, pseudogene, putative |
| Tb927.4.4420 | 9.61 | 1.10E-03 | adenylyl cyclase, pseudogene, putative |
| Tb07.30D13.110 | 9.47 | 2.32E-07 | hypothetical protein, conserved (pseudogene) |
| Tb927.5.4010 | 9.45 | 1.10E-09 | Enriched in surface-labeled proteome protein 4 |
| Tb927.7.390 | 9.32 | 2.30E-05 | hypothetical protein, conserved |
| Tb2.NT.8 | 9.27 | 6.77E-06 | lncRNA, putative |
| Tb927.5.310 | 9.26 | 7.17E-11 | invariant surface glycoprotein, putative |
| KS17gene_6111a | 9.22 | 2.60E-03 | lncRNA, putative |
| Tb927.7.2970 | 9.17 | 7.05E-07 | ATP-dependent DEAD/H RNA helicase, putative |
| Tb927.2.6000 | 9.10 | 9.78E-09 | glycosylphosphatidylinositol-specific phospholipase C |
| KS17gene_372a | 9.06 | 5.29E-05 | lncRNA, putative |
| Tb927.4.4470 | 9.05 | 3.86E-09 | adenylyl cyclase |
| KS17gene_3091a | 9.04 | 1.86E-02 | lncRNA, putative |
| KS17gene_371a | 9.00 | 3.03E-03 | lncRNA, putative |
| nt_1759.1 | 8.88 | 3.77E-04 | lncRNA, putative |
| Tb927.6.3550 | 8.88 | 3.60E-03 | phospholipid-translocating P-type ATPase (flippase), putative |
| Tb927.2.2015 | 8.80 | 6.93E-06 | hypothetical protein |
| Tb927.11.15855 | 8.77 | 9.78E-09 | hypothetical protein |
| Tb927.2.1160 | 8.77 | 3.56E-04 | retrotransposon hot spot protein 2 (RHS2), interrupted, degenerate |
| Tb927.10.8480 | 8.68 | 8.11E-14 | glucose transporter, putative |
| Tb927.2.1200 | 8.59 | 3.25E-09 | DNA-directed RNA polymerase III subunit 2, pseudogene, putative |
| nt_445.1 | 8.57 | 2.40E-09 | lncRNA, putative |
| Tb927.10.1040 | 8.52 | 1.08E-13 | serine peptidase, Clan SC, Family S10 |
| Tb927.4.4440 | 8.44 | 1.91E-10 | adenylyl cyclase |
| nt_256.1 | 8.23 | 7.29E-04 | lncRNA, putative |
| Tb927.8.8030 | 8.21 | 1.89E-11 | emp24/gp25L/p24 family/GOLD, putative |
| nt_430.1 | 8.20 | 2.21E-04 | lncRNA, putative |
| KS17gene_4362a | 8.19 | 4.23E-03 | lncRNA, putative |
| Tb927.7.2030 | 8.17 | 5.85E-03 | retrotransposon hot spot protein 7 (RHS7), putative |
| Tb927.10.2400 | 8.15 | 1.85E-05 | hypothetical protein |
| Tb927.3.5660 | 8.13 | 1.18E-06 | UDP-GlcNAc:alpha3-D-mannoside beta-1,2-N-acetylglucosaminyltransferase I |
| Tb927.2.1250 | 7.98 | 5.14E-06 | hypothetical protein |
| Tb927.7.3830 | 7.97 | 2.41E-05 | kinesin K39, putative |
| KS17gene_1169a | 7.96 | 6.44E-03 | lncRNA, putative |
| Tb927.8.7900 | 7.96 | 5.57E-03 | receptor-type adenylate cyclase GRESAG 4, putative |
| Tb927.5.350 | 7.84 | 2.18E-04 | 75 kDa invariant surface glycoprotein, putative |
| Tb927.2.950 | 7.82 | 1.64E-04 | paraflagellar rod component, putative |
| Tb927.8.6720 | 7.80 | 4.06E-08 | hypothetical protein, conserved |

| | | | |
|---|---|---|---|
| KS17gene_1443a | 7.79 | 5.28E-03 | lncRNA, putative |
| Tb927.1.350 | 7.79 | 1.63E-08 | retrotransposon hot spot protein (RHS, pseudogene), putative |
| Tb927.11.12140 | 7.77 | 6.45E-03 | hypothetical protein |
| Tb927.10.6720 | 7.76 | 7.05E-09 | Plasma-membrane choline transporter, putative |
| Tb927.2.270 | 7.73 | 6.64E-03 | retrotransposon hot spot protein 3 (RHS3), frameshift |
| KS17gene_4607a | 7.72 | 1.56E-04 | lncRNA, putative |
| Tb927.4.3380 | 7.71 | 1.70E-03 | myosin IB heavy chain, putative |
| Tb927.9.15660 | 7.62 | 8.40E-08 | procyclic-enriched flagellar receptor adenylate cyclase 6 |
| Tb1.NT.24 | 7.60 | 3.68E-07 | lncRNA, putative |
| KS17gene_3141a | 7.59 | 4.10E-04 | lncRNA, putative |
| KS17gene_4605a | 7.55 | 1.59E-07 | lncRNA, putative |
| KS17gene_1328a | 7.54 | 2.26E-09 | lncRNA, putative |
| Tb927.10.1050 | 7.52 | 2.22E-08 | serine peptidase, Clan SC, Family S10 |
| Tb927.11.6120 | 7.51 | 7.57E-03 | ABC transporter, putative |
| Tb927.11.17840 | 7.45 | 1.69E-03 | retrotransposon hot spot protein (RHS), degenerate |
| Tb927.6.5160 | 7.44 | 8.47E-06 | retrotransposon hot spot protein 3 (RHS3), point mutation |
| Tb927.6.360 | 7.40 | 3.67E-04 | UDP-Gal or UDP-GlcNAc-dependent glycosyltransferase (pseudogene), putative |
| KS17gene_2695a | 7.35 | 3.93E-03 | lncRNA, putative |
| Tb07.30D13.130 | 7.28 | 8.06E-06 | hypothetical protein, conserved (pseudogene) |
| Tb927.10.6420 | 7.27 | 1.07E-02 | hypothetical protein |
| Tb927.2.1310 | 7.26 | 4.91E-04 | leucine-rich repeat protein (LRRP, pseudogene), putative |
| Tb927.2.1120 | 7.25 | 7.20E-06 | retrotransposon hot spot protein 4 (RHS4), point mutation |
| Tb927.11.15850 | 7.25 | 2.00E-09 | kinteoplast poly(A) polymerase complex 1 subunit |
| Tb927.1.480 | 7.09 | 2.20E-05 | leucine-rich repeat protein (LRRP), putative |
| Tb927.7.1130 | 7.08 | 1.87E-11 | trypanothione/tryparedoxin dependent peroxidase 2 |
| Tb927.9.18140 | 7.01 | 3.91E-10 | variant surface glycoprotein (VSG, pseudogene), putative |
| Tb927.2.300 | 6.98 | 4.22E-10 | DNA-directed RNA polymerase, pseudogene, putative |
| Tb927.9.13070 | 6.96 | 8.02E-15 | Heat shock factor binding 1 domain-containing protein |
| Tb927.10.6740 | 6.92 | 6.19E-11 | Plasma-membrane choline transporter, putative |
| KS17gene_3751a | 6.88 | 2.18E-05 | lncRNA, putative |
| KS17gene_3345a | 6.83 | 4.05E-03 | lncRNA, putative |
| Tb927.1.60 | 6.79 | 1.14E-12 | RNA polymerase (pseudogene), putative |
| KS17gene_5972a | 6.78 | 3.56E-03 | lncRNA, putative |
| Tb927.11.11600 | 6.73 | 1.74E-04 | hypothetical protein, conserved |
| Tb927.11.1520 | 6.72 | 1.75E-04 | expression site-associated gene 3 (ESAG3) protein, putative |
| Tb927.6.440 | 6.71 | 7.06E-07 | haptoglobin-hemoglobin receptor |
| Tb927.5.630 | 6.61 | 2.64E-13 | acidic phosphatase, putative |
| Tb927.2.5330 | 6.59 | 4.84E-03 | hypothetical protein, conserved |
| KS17gene_4164a | 6.56 | 6.01E-08 | lncRNA, putative |
| Tb927.6.160 | 6.56 | 3.09E-08 | retrotransposon hot spot protein 1 (RHS1), putative |
| Tb927.8.6730 | 6.48 | 1.07E-08 | Enriched in surface-labeled proteome protein 24 |
| Tb927_08_v4.snoRNA.0044 | 6.48 | 3.95E-06 | H/ACA-like snoRNA |
| Tb927.11.15870 | 6.47 | 2.08E-09 | hypothetical protein, conserved |
| Tb927.6.3470 | 6.45 | 1.00E-04 | hypothetical protein, conserved |
| Tb927.2.460 | 6.42 | 3.70E-10 | DNA-directed RNA polymerase, pseudogene, putative |
| Tb927.11.4770 | 6.38 | 1.79E-09 | retrotransposon hot spot protein (RHS, pseudogene), putative |
| KS17gene_4419a | 6.35 | 2.63E-11 | lncRNA, putative |
| nt_451.3 | 6.34 | 2.03E-07 | lncRNA, putative |
| Tb927.9.13650 | 6.31 | 1.32E-12 | ADP-ribosylation factor, putative |
| Tb927.4.100 | 6.28 | 1.34E-10 | retrotransposon hot spot protein 1 (RHS1), interrupted |
| Tb927.1.70 | 6.25 | 1.69E-02 | retrotransposon hot spot protein 4 (RHS4), putative |
| Tb927.7.300 | 6.22 | 1.18E-10 | UDP-Gal or UDP-GlcNAc-dependent glycosyltransferase, putative |
| Tb927.9.15930 | 6.22 | 9.30E-08 | small GTPase, putative |
| Tb927.4.180 | 6.16 | 4.87E-07 | hypothetical protein |
| KS17gene_1185a | 6.15 | 1.13E-04 | lncRNA, putative |
| KS17gene_6076a | 6.11 | 1.13E-07 | lncRNA, putative |
| Tb927.10.16190 | 6.09 | 8.28E-06 | procyclic-enriched flagellar receptor adenylate cyclase 2 |
| KS17gene_6855a | 6.08 | 2.41E-03 | lncRNA, putative |
| KS17gene_3712a | 6.06 | 1.96E-04 | lncRNA, putative |
| Tb11.NT.27 | 6.02 | 9.12E-04 | lncRNA, putative |
| Tb927.10.70 | 6.01 | 3.93E-03 | retrotransposon hot spot (RHS), putative, (fragment) |
| KS17gene_6769a | 5.87 | 2.14E-03 | lncRNA, putative |

| | | | |
|---|---|---|---|
| Tb927.11.5910 | 5.83 | 1.95E-05 | WASH complex subunit 7, N-terminal/WASH complex subunit 7/WASH complex subunit 7, C-terminal, putative |
| Tb927.1.380 | 5.73 | 1.80E-08 | Protein of unknown function (DUF1181), putative |
| KS17gene_2586a | 5.71 | 2.52E-03 | lncRNA, putative |
| KS17gene_7754a | 5.70 | 2.56E-03 | lncRNA, putative |
| Tb927.9.8950 | 5.70 | 1.41E-11 | metallo- peptidase, Clan M- Family M48 |
| Tb927.9.11480 | 5.69 | 6.56E-11 | Enriched in surface-labeled proteome protein 9 |
| Tb927.11.5210 | 5.66 | 1.77E-03 | hypothetical protein, conserved |
| Tb927.10.3970 | 5.65 | 4.80E-12 | hypothetical protein, conserved |
| Tb927.7.6590 | 5.64 | 1.26E-04 | hypothetical protein, conserved |
| Tb927.10.14160 | 5.64 | 6.43E-10 | Aquaglyceroporin 3 |
| Tb927.3.5560 | 5.64 | 3.15E-06 | hypothetical protein, conserved |
| Tb927.10.14140 | 5.63 | 1.24E-13 | pyruvate kinase 1 |
| Tb927.1.120 | 5.59 | 2.03E-13 | retrotransposon hot spot protein 4 (RHS4), putative |
| Tb927.7.470 | 5.51 | 2.57E-07 | Enriched in surface-labeled proteome protein 14 |
| Tb927.10.8230 | 5.51 | 1.69E-13 | protein disulfide isomerase 2 |
| KS17gene_8043a | 5.49 | 1.28E-02 | lncRNA, putative |
| Tb927.4.4580 | 5.46 | 1.68E-03 | hypothetical protein, conserved |
| Tb927.9.13380 | 5.45 | 8.10E-10 | Autophagy-related protein 24 |
| Tb927.7.6490 | 5.44 | 2.59E-06 | hypothetical protein, conserved |
| Tb927.2.510 | 5.43 | 4.96E-04 | retrotransposon hot spot protein 4 (RHS4), putative |
| Tb927.6.300 | 5.42 | 1.18E-02 | receptor-type adenylate cyclase GRESAG 4, putative |
| Tb927.8.3900 | 5.41 | 1.34E-07 | hypothetical protein, conserved |
| KS17gene_4037a | 5.39 | 3.82E-05 | lncRNA, putative |
| Tb927_08_v4.snoRNA.0043 | 5.38 | 6.19E-03 | H/ACA-like snoRNA |
| KS17gene_2098a | 5.37 | 2.03E-04 | lncRNA, putative |
| Tb927.1.3670 | 5.36 | 7.41E-05 | expression site-associated gene 8 (ESAG8) protein, putative |
| Tb927.2.5350 | 5.34 | 3.66E-03 | hypothetical protein, conserved |
| KS17gene_846a | 5.33 | 1.34E-02 | lncRNA, putative |
| Tb927.2.170 | 5.32 | 1.62E-11 | leucine-rich repeat protein 1 (LRRP1), putative |
| Tb927.11.11965 | 5.31 | 3.37E-12 | cytoskeleton-associated protein, putative |
| Tb927.2.1080 | 5.30 | 1.64E-11 | retrotransposon hot spot protein 5 (RHS5), putative |
| KS17gene_5447a | 5.27 | 1.70E-03 | lncRNA, putative |
| Tb927.4.2070 | 5.23 | 8.14E-07 | antigenic protein, putative |
| Tb927.2.900 | 5.16 | 2.42E-08 | hypothetical protein |
| Tb927.10.9710 | 5.15 | 1.99E-03 | hypothetical protein |
| KS17gene_3132a | 5.14 | 1.40E-02 | lncRNA, putative |
| Tb927.7.1120 | 5.14 | 6.57E-10 | trypanothione/tryparedoxin dependent peroxidase 1, cytosolic |
| Tb927.11.15490 | 5.08 | 2.89E-05 | Tb-291 membrane associated protein, putative |
| Tb927.7.6860 | 5.08 | 3.00E-14 | expression site-associated gene 5 (ESAG5) protein, putative |
| nt_17.3 | 5.06 | 2.70E-04 | lncRNA, putative |
| Tb927.4.3980 | 5.01 | 1.02E-06 | chaperone protein DnaJ, putative |
| nt_6689.1 | 5.00 | 7.38E-03 | lncRNA, putative |
| KS17gene_6292a | 4.96 | 2.17E-02 | lncRNA, putative |
| Tb927.11.11980 | 4.96 | 4.14E-12 | cytoskeleton-associated protein 15 |
| Tb927.2.380 | 4.95 | 2.28E-07 | retrotransposon hot spot protein 2 (RHS2), putative |
| Tb927.2.160 | 4.91 | 5.86E-03 | Protein of unknown function (DUF1181), putative |
| KS17gene_5090a | 4.90 | 3.26E-07 | lncRNA, putative |
| Tb927.7.6070 | 4.90 | 1.04E-04 | receptor-type adenylate cyclase GRESAG 4, putative |
| KS17gene_3810a | 4.90 | 5.80E-04 | lncRNA, putative |
| Tb927.10.8450 | 4.90 | 2.22E-02 | glucose transporter 1E |
| KS17gene_7369a | 4.89 | 9.04E-08 | lncRNA, putative |
| Tb927.2.560 | 4.88 | 5.22E-10 | retrotransposon hot spot protein 4 (RHS4), putative |
| KS17gene_4365a | 4.85 | 6.01E-03 | lncRNA, putative |
| KS17gene_329a | 4.83 | 3.52E-03 | lncRNA, putative |
| Tb927.9.2520 | 4.82 | 6.77E-07 | microtubule-associated protein |
| Tb927.10.4180 | 4.81 | 6.82E-11 | TFIIF-stimulated CTD phosphatase, putative |
| KS17gene_6325a | 4.76 | 1.84E-08 | lncRNA, putative |
| Tb927.6.560 | 4.75 | 7.87E-12 | cysteine peptidase C (CPC) |
| Tb927.5.610 | 4.74 | 7.44E-10 | acidic phosphatase, putative |
| Tb927.10.1770 | 4.73 | 1.02E-04 | hypothetical protein |
| Tb927.6.130 | 4.73 | 1.54E-05 | hypothetical leucine-rich repeat protein 1 (LRRP1) |

| | | | |
|---|---|---|---|
| Tb927.10.10490 | 4.70 | 5.47E-12 | histone H2B, putative |
| Tb927.11.7080 | 4.69 | 5.31E-09 | acidocalcisomal pyrophosphatase |
| Tb927.9.15700 | 4.68 | 2.35E-02 | variant surface glycoprotein (VSG)-related, putative |
| Tb927.2.1344 | 4.68 | 2.10E-02 | retrotransposon hot spot protein 1 (RHS1), interrupted |
| KS17gene_4884a | 4.67 | 4.43E-03 | lncRNA, putative |
| Tb927.7.3840 | 4.66 | 2.43E-02 | kinesin-like protein, fragment, putative |
| KS17gene_5952a | 4.66 | 4.56E-06 | lncRNA, putative |
| Tb927.5.292b | 4.64 | 1.48E-03 | hypothetical protein |
| Tb927.11.1230 | 4.59 | 7.98E-05 | hypothetical protein, conserved |
| Tb927.3.5180 | 4.59 | 1.72E-11 | ADF/Cofilin |
| Tb927.5.840 | 4.56 | 4.52E-03 | Nucleolar protein 111 |
| Tb927.11.13040 | 4.54 | 1.14E-06 | calmodulin |
| Tb927.7.4260 | 4.50 | 3.12E-09 | Enriched in surface-labeled proteome protein 13 |
| KS17gene_4614a | 4.48 | 3.63E-08 | lncRNA, putative |
| Tb927.3.5780 | 4.47 | 3.25E-05 | hypothetical protein |
| KS17gene_7740a | 4.45 | 9.88E-03 | lncRNA, putative |
| Tb927.1.4970 | 4.44 | 1.11E-02 | hypothetical protein |
| Tb927.6.150 | 4.42 | 2.13E-03 | retrotransposon hot spot protein 3 (RHS3), frameshift |
| Tb927.2.6150 | 4.40 | 2.05E-04 | adenosine transporter 2 |
| nt_255.1 | 4.39 | 5.18E-03 | lncRNA, putative |
| Tb6.NT.40 | 4.36 | 1.11E-03 | lncRNA, putative |
| Tb927.6.3940 | 4.36 | 1.85E-07 | Autophagy-related protein 27, putative |
| KS17gene_5976a | 4.35 | 2.93E-05 | lncRNA, putative |
| Tb927.11.11320 | 4.34 | 9.26E-05 | hypothetical protein, conserved |
| Tb927.8.2260 | 4.34 | 3.01E-07 | Present in the outer mitochondrial membrane proteome 39-2 |
| Tb927.5.4590 | 4.29 | 2.96E-03 | small GTP-binding rab protein, pseudogene, putative |
| Tb927.5.2410 | 4.29 | 1.01E-05 | kinesin, putative |
| Tb927.8.5100 | 4.28 | 1.07E-03 | hypothetical protein, conserved |
| Tb927.8.3890 | 4.27 | 5.95E-08 | hypothetical protein, conserved |
| Tb927.11.12800 | 4.27 | 1.08E-05 | ribonucleoside-diphosphate reductase small chain |
| KS17gene_213a | 4.26 | 3.32E-04 | lncRNA, putative |
| nt_417.2 | 4.21 | 6.96E-05 | lncRNA, putative |
| Tb927.7.2650 | 4.21 | 6.24E-10 | Cytoskeleton associated protein 51V |
| Tb927.5.4380 | 4.20 | 7.96E-04 | Kinetoplastid-specific Protein Phosphatase 1 |
| Tb927.8.4060 | 4.19 | 5.37E-09 | Flagellum adhesion protein 2, putative |
| Tb927.3.3450 | 4.16 | 1.51E-12 | ADP-ribosylation factor-like protein 3, putative |
| Tb927.1.270 | 4.15 | 2.23E-05 | hypothetical protein |
| Tb927.10.8350 | 4.13 | 1.87E-02 | hypothetical protein |
| Tb927.7.480 | 4.13 | 3.12E-04 | hypothetical protein, conserved |
| Tb927_09_v4.snoRNA.0027 | 4.11 | 7.02E-03 | C/D snoRNA |
| Tb927.1.4710 | 4.11 | 1.65E-03 | hypothetical protein, conserved |
| Tb927.1.3860 | 4.08 | 2.91E-04 | hypothetical protein, conserved |
| KS17gene_4622a | 4.06 | 1.94E-10 | lncRNA, putative |
| Tb927.5.360 | 4.05 | 4.00E-12 | 75 kDa invariant surface glycoprotein |
| KS17gene_1897a | 4.05 | 1.88E-04 | lncRNA, putative |
| Tb927.9.6570 | 4.03 | 3.34E-03 | hypothetical protein, conserved |
| Tb927.11.5370 | 4.02 | 9.19E-07 | hypothetical protein, conserved |
| Tb927.4.1200 | 4.02 | 1.87E-03 | expression site-associated gene 1 (ESAG1) protein, putative |

**Supplementary Table S7.** Differential gene expression. Genes up-regulated in procyclic-stage trypanosomes.

| **gene-ID** | **fold change (PC/BS)** | **adjusted *p*-value** | **gene product** |
|---|---|---|---|
| Tb927.10.10270 | 61.83 | 4.82E-24 | hypothetical protein |
| Tb927.10.10260 | 44.01 | 7.95E-30 | EP1 procyclin |
| Tb927.10.10250 | 33.52 | 2.17E-21 | EP2 procyclin |
| KS17gene_6791a | 32.87 | 1.50E-10 | lncRNA, putative |
| Tb927.3.590 | 32.66 | 1.65E-17 | adenosine transporter, putative |
| Tb927.6.450 | 28.44 | 7.21E-23 | procyclin PARP A |
| Tb927.6.520 | 26.59 | 6.68E-19 | EP3-2 procyclin |
| Tb927.5.2260 | 26.29 | 8.65E-24 | conserved protein |
| nt_6401.1 | 22.24 | 9.88E-07 | lncRNA, putative |
| Tb927.4.4000 | 22.15 | 9.28E-12 | amino acid transporter, putative |
| Tb927.7.3020 | 20.76 | 7.21E-23 | Nitroreductase family, putative |
| nt_3670.2 | 19.73 | 8.15E-06 | lncRNA, putative |
| Tb927.7.2980 | 18.85 | 2.05E-23 | Nitroreductase family, putative |
| Tb927.7.430 | 18.63 | 3.94E-07 | hypothetical protein, conserved |
| Tb927.7.440 | 18.60 | 8.98E-07 | hypothetical protein, conserved |
| Tb927.10.10240 | 18.21 | 1.78E-09 | procyclin-associated gene 1 (PAG1) protein |
| Tb927.10.15410 | 16.90 | 6.09E-19 | glycosomal malate dehydrogenase |
| Tb927.11.7500 | 16.77 | 1.75E-20 | Protein of unknown function (DUF423), putative |
| Tb927.5.4020 | 16.41 | 2.72E-12 | hypothetical protein |
| Tb927.6.3490 | 16.06 | 3.23E-16 | zinc finger protein 1 |
| Tb927.7.340 | 15.85 | 3.05E-05 | Alpha/beta hydrolase family, putative |
| Tb927.6.480 | 15.55 | 8.03E-12 | surface protein EP3-2 procyclin precursor |
| Tb927.5.990 | 14.80 | 5.74E-10 | hypothetical protein, conserved |
| Tb927.10.8490 | 13.86 | 4.26E-19 | glucose transporter, putative |
| Tb927.7.6883 | 13.77 | 3.64E-11 | 28S alpha ribosomal RNA |
| Tb927.11.6280 | 13.04 | 2.56E-16 | pyruvate phosphate dikinase |
| Tb927.11.1710 | 12.57 | 2.17E-21 | guide RNA-binding protein of 21 kDa |
| Tb927.11.2410 | 12.52 | 1.65E-17 | Flabarin, putative |
| Tb927.2.4610 | 12.29 | 1.71E-18 | branched-chain amino acid aminotransferase, putative |
| Tb927.7.6180 | 12.21 | 2.59E-04 | hypothetical protein, conserved |
| Tb927.6.3880 | 12.13 | 4.11E-05 | hypothetical protein, conserved |
| Tb927.8.480 | 12.00 | 1.44E-05 | phosphatidic acid phosphatase protein, putative |
| Tb927.6.970 | 11.94 | 1.69E-16 | cysteine peptidase, Clan CA, family C1, Cathepsin L-like |
| Tb927.9.15550 | 11.83 | 4.49E-06 | BARP protein |
| Tb927.11.7490 | 11.64 | 2.84E-18 | Protein of unknown function (DUF423), putative |
| Tb927.11.4700 | 11.37 | 2.17E-21 | prostaglandin f synthase |
| Tb927.8.8300 | 10.88 | 4.80E-06 | amino acid transporter, putative |
| Tb927.10.15090 | 10.73 | 1.77E-16 | hypothetical protein, conserved |
| Tb927.1.710 | 10.57 | 2.61E-17 | phosphoglycerate kinase |
| Tb927.8.2540 | 10.16 | 2.27E-18 | 3-ketoacyl-CoA thiolase |
| Tb927.3.3449 | 10.08 | 1.42E-05 | 28S alpha ribosomal RNA |
| Tb927.10.9080 | 9.85 | 7.23E-14 | pteridine transporter, putative |
| Tb927.11.3730 | 9.82 | 1.08E-06 | leucyl-tRNA synthetase, putative |
| Tb927.7.2210 | 9.75 | 4.89E-05 | hypothetical protein, conserved |
| Tb927.6.1010 | 9.69 | 9.81E-15 | cysteine peptidase, Clan CA, family C1, Cathepsin L-like |
| Tb927.10.4280 | 9.47 | 4.44E-15 | ubiquinol-cytochrome c reductase complex 14 kda protein |
| Tb927.11.9980 | 9.37 | 3.01E-07 | 2-oxoglutarate dehydrogenase E1 component, putative |
| Tb927.9.7470 | 9.28 | 9.60E-10 | purine nucleoside transporter |
| Tb927.10.13430 | 9.20 | 2.03E-05 | citrate synthase, putative |
| Tb927.7.5970 | 8.62 | 4.98E-16 | protein associated with differentiation 5, putative |
| Tb927.11.5440 | 8.46 | 1.45E-12 | NADP-dependent malic enzyme, cytosolic |
| Tb927.10.8750 | 8.39 | 5.98E-04 | GTPase activating protein, putative |
| Tb927.6.500 | 8.31 | 6.12E-10 | gene related to expression site-associated gene 2 (GRESAG2), putative |
| Tb927.2.4210 | 7.98 | 1.13E-17 | Phosphoenolpyruvate carboxykinase [ATP], glycosomal |
| Tb927.10.15900 | 7.85 | 8.43E-09 | hypothetical protein, conserved |
| Tb927.11.8990 | 7.79 | 8.27E-08 | cation transporter, putative |
| Tb927.11.15550 | 7.75 | 3.93E-08 | NADH-cytochrome b5 reductase, putative |
| Tb927.8.6750 | 7.68 | 1.09E-13 | translationally controlled tumor protein (TCTP), putative |
| Tb927.8.1550 | 7.66 | 1.77E-03 | paraflagellar rod component, putative |
| Tb927.10.2190 | 7.54 | 9.99E-04 | Protein of unknown function (DUF667), putative |
| Tb927.5.2160 | 7.48 | 7.46E-12 | conserved protein |

| | | | |
|---|---|---|---|
| Tb927.10.3210 | 7.43 | 1.34E-12 | delta-1-pyrroline-5-carboxylate dehydrogenase, putative |
| Tb927.10.2560 | 7.13 | 4.82E-15 | mitochondrial malate dehydrogenase |
| Tb927.10.9350 | 7.12 | 1.06E-04 | hypothetical protein, conserved |
| Tb927.10.5760 | 7.10 | 1.29E-04 | adenylate kinase, putative |
| Tb927.6.530 | 7.09 | 1.17E-11 | procyclin associated gene 3 (PAG3) protein |
| Tb927.10.8500 | 7.05 | 1.42E-10 | glucose transporter, putative |
| Tb927.11.3610 | 6.96 | 2.43E-13 | nucleobase/nucleoside transporter 8.1 |
| Tb927.3.1840 | 6.95 | 4.09E-06 | 3-oxo-5-alpha-steroid 4-dehydrogenase, putative |
| Tb927.6.760 | 6.94 | 2.02E-05 | receptor-type adenylate cyclase GRESAG 4, putative |
| Tb927.11.11360 | 6.90 | 9.22E-20 | receptor for activated C kinase 1 |
| Tb927.5.2560 | 6.74 | 2.73E-12 | hypothetical protein, conserved |
| Tb11.02.5400 | 6.70 | 1.17E-11 | cystathionine beta-synthase, putative |
| Tb927.7.4070 | 6.68 | 1.62E-14 | cysteine peptidase, Clan CA, family C2, putative |
| Tb927.3.3432 | 6.59 | 3.79E-06 | 28S alpha ribosomal RNA |
| Tb927.11.rRNA_1 | 6.56 | 1.45E-12 | 5.8S ribosomal RNA |
| Tb927.1.3800 | 6.51 | 8.41E-04 | Present in the outer mitochondrial membrane proteome 18 |
| Tb927.10.2350 | 6.48 | 2.24E-06 | pyruvate dehydrogenase complex E3 binding protein, putative |
| Tb927.4.3950 | 6.35 | 4.35E-09 | cytoskeleton-associated protein CAP5.5, putative |
| Tb927.3.3441 | 6.35 | 1.47E-05 | 28S alpha ribosomal RNA |
| Tb927.10.12780 | 6.32 | 9.89E-07 | Zinc finger CCCH domain-containing protein 37 |
| Tb927.1.2210 | 6.26 | 2.61E-07 | nucleosome assembly protein (NAP), putative |
| Tb927.1.4630 | 6.18 | 3.58E-03 | cyclin-like F-box protein 1E |
| Tb927.9.7920 | 6.14 | 2.31E-03 | hypothetical protein, conserved |
| Tb927.9.15540 | 6.04 | 3.11E-08 | BARP protein |
| Tb927.7.1320 | 5.97 | 2.40E-17 | 10 kDa heat shock protein, putative |
| Tb927.10.4030 | 5.95 | 1.14E-03 | hypothetical protein |
| Tb927.4.1360 | 5.92 | 1.97E-09 | Glucose-6-phosphate 1-epimerase, putative |
| Tb927.6.2070 | 5.92 | 5.85E-05 | Mitoribosomal SSU assembly factor 28 |
| Tb927.6.200 | 5.91 | 3.24E-06 | receptor-type adenylate cyclase GRESAG 4, putative |
| Tb927.8.6170 | 5.87 | 1.86E-12 | transketolase, putative |
| Tb927.10.10770 | 5.86 | 2.20E-11 | Generative cell specific 1 protein, putative |
| Tb927.10.8530 | 5.84 | 1.36E-10 | glucose transporter 2A |
| Tb927.10.6860 | 5.83 | 5.04E-04 | endonuclease v |
| Tb927.2.1443 | 5.75 | 1.64E-12 | 5.8S ribosomal RNA |
| Tb927.11.4760 | 5.72 | 1.30E-07 | hypothetical protein |
| Tb927.7.4390 | 5.70 | 1.87E-12 | threonine synthase, putative |
| Tb927.9.6760 | 5.62 | 2.27E-03 | hypothetical protein, conserved |
| Tb927.10.470 | 5.51 | 5.43E-09 | choline dehydrogenase, putative |
| Tb927.8.4720 | 5.49 | 1.16E-12 | amino acid transporter, putative |
| Tb927.8.7670 | 5.42 | 3.68E-09 | amino acid transporter, putative |
| Tb927.1.2820 | 5.35 | 8.34E-13 | pteridine transporter, putative |
| Tb927.11.3270 | 5.34 | 3.42E-04 | squalene monooxygenase, putative |
| Tb927.10.11220 | 5.32 | 7.51E-14 | procyclic form surface phosphoprotein |
| Tb927.2.3920 | 5.31 | 4.84E-08 | Complex 1 protein (LYR family), putative |
| Tb927.7.5400 | 5.28 | 4.06E-05 | hypothetical protein, conserved |
| Tb927.11.16730 | 5.25 | 2.02E-08 | dihydrolipoyl dehydrogenase |
| Tb927.11.12490 | 5.25 | 1.86E-03 | hypothetical protein, conserved |
| Tb927.6.4790 | 5.24 | 1.89E-10 | hypothetical protein, conserved |
| Tb927.9.1520 | 5.22 | 1.12E-13 | hypothetical protein, conserved |
| Tb927.1.2880 | 5.20 | 3.53E-11 | pteridine transporter, putative |
| Tb927.3.3890 | 5.10 | 1.26E-05 | hypothetical protein, conserved |
| Tb927.7.6850 | 5.08 | 2.19E-08 | trans-sialidase |
| Tb927.1.2470 | 5.03 | 9.81E-15 | histone H3, putative |
| KS17gene_1079a | 5.00 | 1.01E-09 | lncRNA, putative |
| Tb927.9.2320 | 4.96 | 2.14E-05 | methyltransferase domain containing protein, putative |
| Tb927.11.9750 | 4.94 | 1.08E-06 | Protein of unknown function (DUF498/DUF598), putative |
| Tb927.11.14000 | 4.94 | 3.94E-16 | nuclear RNA binding domain 1 |
| Tb927.7.5990 | 4.94 | 8.54E-08 | protein associated with differentiation 7, putative |
| Tb927.6.610 | 4.91 | 3.27E-04 | kinetoplast ribosomal PPR-repeat containing protein 18 |
| Tb927.10.10000 | 4.91 | 8.34E-13 | hypothetical protein, conserved |
| Tb927.9.7980 | 4.91 | 1.45E-12 | hypothetical protein, conserved |
| Tb927.9.2450 | 4.88 | 2.60E-03 | electon transport protein SCO1/SCO2, putative |
| Tb927.4.4620 | 4.80 | 4.94E-13 | cytochrome oxidase subunit VIII |
| Tb927.10.690 | 4.78 | 4.20E-06 | palmitoyl acyltransferase 3, putative |
| Tb927.2.1953 | 4.77 | 2.73E-06 | 28S alpha ribosomal RNA |
| Tb927.9.15580 | 4.73 | 8.06E-09 | BARP protein |
| Tb927.10.6200 | 4.72 | 9.04E-09 | hypothetical protein, conserved |
| Tb927.6.3800 | 4.69 | 1.73E-08 | heat shock 70 kDa protein, mitochondrial precursor, putative |
| Tb927.1.2400 | 4.67 | 2.68E-16 | alpha tubulin |

| | | | |
|---|---|---|---|
| Tb927.10.15950 | 4.63 | 2.93E-05 | TATA-box-binding protein |
| Tb927.3.700 | 4.60 | 4.16E-09 | hypothetical protein, conserved |
| Tb927.5.3890 | 4.59 | 8.09E-06 | hypothetical protein, conserved |
| Tb927.8.1640 | 4.59 | 2.05E-13 | MSP-B, putative |
| Tb927.3.3750 | 4.59 | 3.26E-07 | paraflagellar rod component, putative |
| Tb927.8.4010 | 4.53 | 2.24E-06 | Flagellum adhesion protein 1 |
| Tb927.5.440 | 4.50 | 1.06E-09 | trans-sialidase, putative |
| Tb927.11.7260 | 4.48 | 3.06E-07 | hypothetical protein, conserved |
| Tb927.6.5095 | 4.41 | 1.99E-12 | hypothetical protein |
| Tb927.11.12440 | 4.40 | 8.29E-05 | Plus-3 domain/Zinc finger, C3HC4 type (RING finger), putative |
| Tb927.4.3990 | 4.39 | 1.74E-04 | amino acid transporter, putative |
| Tb927.5.1060 | 4.38 | 1.95E-09 | mitochondrial processing peptidase, beta subunit, putative |
| Tb927.3.3423 | 4.35 | 5.71E-05 | 28S alpha ribosomal RNA |
| Tb927.7.5980 | 4.35 | 5.79E-04 | protein associated with differentiation 6, putative |
| Tb927.11.1800 | 4.35 | 3.91E-14 | histone H1 |
| Tb927.9.5890 | 4.33 | 1.75E-11 | solanesyl-diphosphate synthase, putative |
| Tb927.7.1790 | 4.23 | 1.62E-07 | Adenine phosphoribosyltransferase, putative |
| Tb927.9.7620 | 4.14 | 5.51E-16 | 60S ribosomal protein L11, putative |
| Tb927.9.14160 | 4.13 | 9.43E-08 | rieske iron-sulfur protein, mitochondrial precursor |
| Tb927.2.3030 | 4.10 | 8.64E-05 | ATP-dependent Clp protease subunit, heat shock protein 78 (HSP78), putative |
| Tb927.3.2880 | 4.10 | 2.29E-09 | Mitochondrial ATP synthase subunit, putative |
| Tb927.5.930 | 4.07 | 1.20E-08 | NADH-dependent fumarate reductase |
| Tb927.11.13140 | 4.03 | 2.27E-12 | cytochrome oxidase subunit X |
| Tb927.3.3431 | 4.01 | 5.63E-11 | 5.8S ribosomal RNA |
| Tb927.9.7830 | 4.00 | 1.03E-11 | tRNA import complex component, putative |
